# Higher-order chromatin organization defines Progesterone Receptor and PAX2 binding to regulate estradiol-primed endometrial cancer gene expression

**DOI:** 10.1101/739466

**Authors:** Alejandro La Greca, Nicolás Bellora, Francois Le Dily, Rodrigo Jara, Javier Quilez Oliete, José Luis Villanueva, Enrique Vidal, Gabriela Merino, Cristóbal Fresno, Inti Tarifa Rieschle, Griselda Vallejo, Guillermo P. Vicent, Elmer Fernández, Miguel Beato, Patricia Saragüeta

**Affiliations:** Biology and Experimental Medicine Institute, IBYME-CONICET, Buenos Aires, Argentina; Centre for Genomic Regulation (CRG), Barcelona Institute for Science and Technology, Barcelona, Spain; Biodiversity and Environment Investigations Institute (INBIOMA), Bariloche, Argentina; Bioscience Data Mining Group, Córdoba University, Córdoba, Argentina; Universitat Pompeu Fabra (UPF), Barcelona, Spain; Consejo Nacional de Investigaciones Científicas y Técnicas (CONICET), Buenos Aires, Argentina

**Keywords:** steroid receptors, gene regulation, endometrial cancer, ChlPseq, Hi-C, ATACseq, progesterone receptor, estrogen receptor, PAX2

## Abstract

Estrogen (E2) and Progesterone (Pg), via their specific receptors (ER and PR respectively), are major determinants in the development and progression of endometrial malignancies. Here, we have studied how E2 and the synthetic progestin R5020 affect genomic functions in Ishikawa endometrial cancer cells. Using ChIPseq in cells exposed to the corresponding hormones, we identified cell specific binding sites for ER (ERbs) and PR (PRbs), which mostly correspond to independent sites but both adjacent to sites bound by PAX2. Analysis of long-range interactions by Hi-C showed enrichment of regions co-bound by PR and PAX2 inside TADs that contain differentially progestin-regulated genes. These regions, which we call “progestin control regions” (PgCRs), exhibit an open chromatin state prior to the exposure to the hormone. Our observations suggest that endometrial response to progestins in differentiated endometrial tumor cells results in part from binding of PR together with partner transcription factors to PgCRs, compartmentalizing hormone-independent open chromatin.

## Introduction

Progesterone (Pg) is a key regulator in the female reproductive tract, including uterine and mammary gland development (Lydon et al., 1995). Endometrial and breast tissues exhibit significantly different responses to hormones, resulting in very distinctive morphologies and functions. During pregnancy, Pg prepares the uterine epithelium to receive the embryo and initiates the process of differentiation of stromal cells towards their decidual phenotype. In the mammary gland and in coordination with prolactin, Pg stimulates epithelial proliferation and differentiation of alveolar lobes in the mammary gland (Mulac-Jericevic and Conneely, 2004). Unlike Pg, estradiol (E2) is the main proliferative signal in the uterine epithelium and exerts its function through activating estrogen receptor (ER) alpha and beta (ERalpha and *β*, respectively) (Ishiwata et al., 1997; Kayisli et al., 2004).

The physiological role of Pg is mediated by the interaction and consequent activation of isoforms A (PRA) and B (PRB) of the progesterone receptor (PR), which are transcribed from alternate promoters of the gene (Hovland et al., 1998). While PRA is more abundant in stromal endometrial cells, PRB is the most representative isoform in ephitelial cells of endometrium. Steroid hormones exert their transcriptional effects through binding of the steroid receptors (SR) to specific DNA sequences in the promoters or enhancers of target genes known as “hormone response elements” (HRE). Estradiol exposure triggers ER binding to estrogen response elements (ERE) regulating target genes such as *PGR*. Previous work showed E2-dependent upregulation of PR in many different target cells, species and pathological conditions (Graham et al., 1995; Kraus and Katzenellenbogen, 1993). Exposure to progestins triggers binding of PR to PRE. Once bound to their HREs the hormone receptors interact with other transcription factors, co-regulators (Beato et al., 1995), such as the p160 family of co-activators of steroid receptors SRC-1-3, and chromatin remodelling enzymes. This evidence favors tissue specific roles of PR isoforms and their co-regulators orientated towards differential transactivation of target genes.

High levels of PRA and PRB have been described in endometrial hyperplasia (Miyamoto et al., 2004) while low and high-grade endometrial cancers reveal reduced or absent expression of one or both isoforms in epithelia or stroma (Shao, 2013). This PR decrease is often associated with shorter progression-free survival and overal survival rates (Leslie et al., 1997; Miyamoto et al., 2004; Sakaguchi et al., 2004; Jongen et al., 2009; Kreizman-Shefer et al., 2014). The absence of PR gene expression may be attributed to hypermethylation of CpG islands within the promoter or first exon regions of the PR gene or to the presence of associated deacetylated histones. These modifications were reported for endometrial cancer cell lines as well as tumor samples and may be exclusive to PRB (Sasaki et al., 2001; Xiong et al., 2005; Ren et al., 2007). Treatment of such cells with DNA methyltranferase or histone deacetylase inhibitors can restore both PRB expression and its regulation of target genes such as *FOXO1*, p21 (*CDKN1A*), p27 (*CDKN1B*), and cyclin D1 (*CCND1*) (Xiong et al., 2005; Yang et al., 2014). Down-regulation of PR by post-transcriptional mechanisms and through pos-translational modifications of PR may contribute to progesterone resistance in endometrial cancer but have not been extensively explored in the context of endometrial cancer. It is known that oncogenic activation of KRAS, PI3K or AKT and/or loss of functional tumor suppressors such as *PTEN* are common genetic alterations (Hecht and Mutter, 2006), toghether with *ARID1A* (Liang et al., 2012), all of them observed in endometrial cancer. Although there are numerous reports of hormonally regulated enhancers and super-enhancers in mammary cancer cells (see in dbsuperenahncer, http://bioinfo.au.tsinghua.edu.cn/dbsuper/) (Khan and Zhang, 2016; Hnisz et al., 2015), there is a void of information about their presence in endometrial cells.

To better understand the response to progestin in endometrial cancer cells, we have studied the genomic binding of ER and PR, the global gene expression changes and the state of chromatin by ATACseq as well as the genomic interactions by HiC in Ishikawa cells exposed to progestin or estrogen, and also in cells exposed to progestin after a period of estradiol pretreatment. Inside TADs with progestin regulated genes, we identified regions that we named “progestin control regions” (PgCRs) that correlate with the open chromatin compartment independently of hormonal stimuli and include binding sites for the partner transcription factor PAX2.

## Results

### Ishikawa endometrial epithelial cells respond to R5020 through activation of PR, whose levels increase upon exposure to E2

Endometrial epithelial cells respond to ovarian steroid hormones −progesterone (Pg) and estradiol (E2)-, E2 being the main proliferative stimulus and Pg its antagonist. After treating Ishikawa cells with E2 10nM for 48h we observed an increment in number of cells compared to vehicle (OH) (FC 1.78±0.08 v. OH) that was suppressed by addition of R5020 10nM (FC 1.15±0.08 v. OH) (Figure 1A). Treatment with R5020 10nM alone did not induce proliferation on Ishikawa cells (FC 0.77±0.08 v. OH) (Figure 1A). E2-induced cell proliferation was also abrogated by pre-incubation with estrogen receptor (ER) antagonist ICI182780 1*μ*M (ICI 10^−6^M) (FC 1.05±0.05 v. OH) (Supplementary Fig. S1A), but not pre-incubation with PR antagonist RU486 1*μ*M (RU486 10^−6^M) (FC 1.42±0.07 v. OH) (Supplementary Fig. S1B), proving that ER but not PR was directly involved in the proliferative response to E2. Suppression of E2-induced cell proliferation by R5020 was inhibited by pre-incubation with RU486 (FC 1.50±0.06 v. OH), indicating that R5020 effect was mediated by PR in Ishikawa cells (Supplementary Fig. S1B). The effects of E2 and R5020 on proliferation were corroborated by BrdU incorporation and cell cycle phase analysis 18h after hormone exposure (Supplementary Fig. S1C and S1D). E2 increased the number of BrdU positive cells and percentage of cells in S phase compared to untreated control cells and to cell exposed to the vehicle (OH), and these increments were inhibited by R5020. Treatment with the histone deacetylase inhibitor Trichostatin A 250nM (TSA 250nM) was used as negative control for BrdU incorporation and cell cycle progression (Supplementary Fig. S1C).

**Figure 1.**
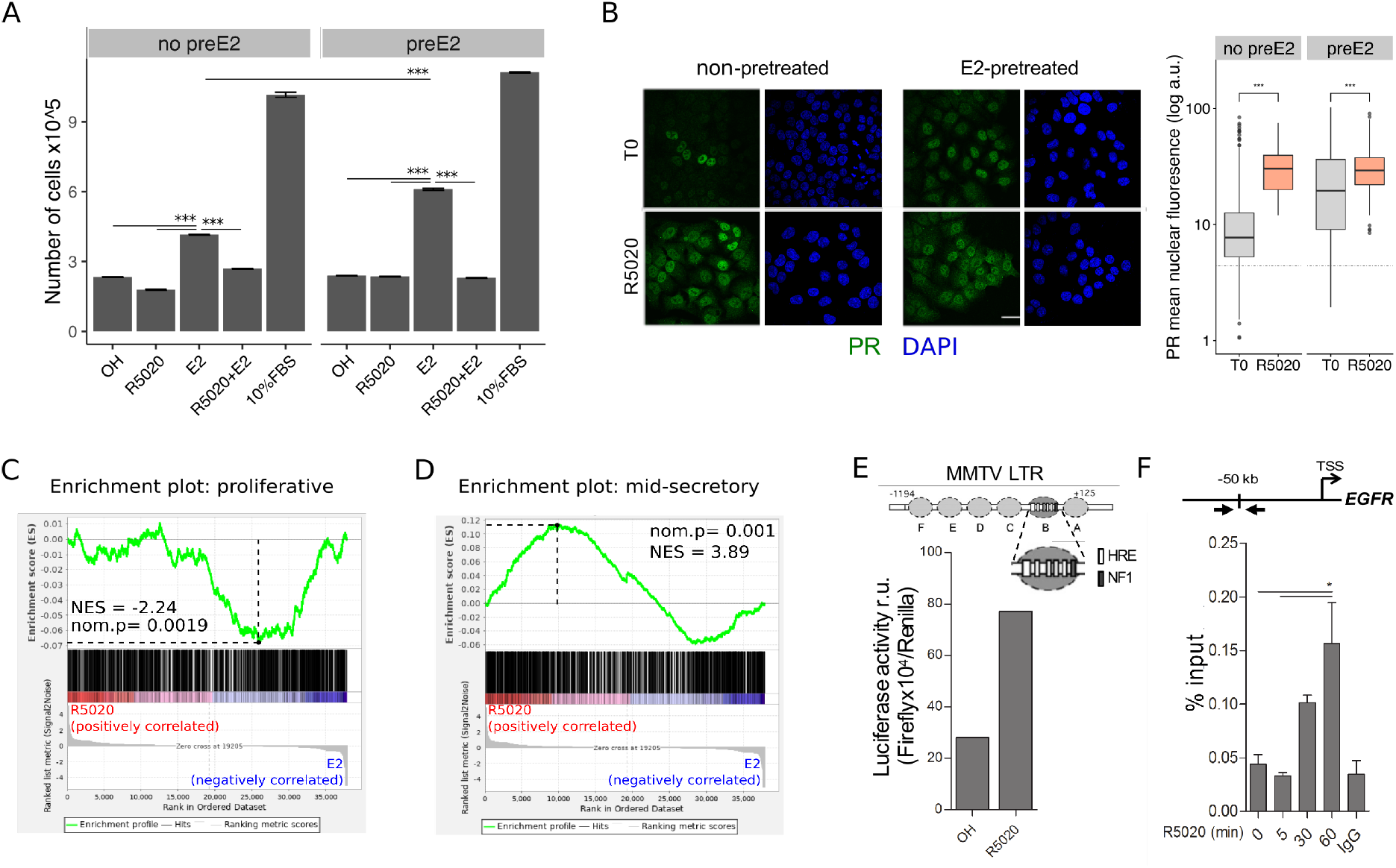
R5020 inhibits E2-induced Ishikawa cell proliferation through an active PR that is capable of transactivating an exogenous MMTV promoter sequence and an endogenous enhancer sequence located 50kb upstream of EGFR gene. (A) Proliferation of Ishikawa cells either pretreated with E2 10nM for 12h (preE2) or not (no preE2) and later treated with vehicle (OH), E2 10nM (E2), R5020 10nM (R5020), E2 combined with R5020 (E2+R5020) and FBS (10%FBS), expressed as mean number of cells ± SE of three independent experiments. (***) p<0.001. (B) Immunofluorescence of PR in untreated (T0; top left), 60min R5020-treated (R5020; bottom left), 12h E2-pretreated (top right) and 12h E2-pretreated 60min R5020-treated (bottom right) Ishikawa cells. Scale bar is equivalent to 30*μ*m. Mean nuclear signal of PR for every cell in all images was determined and shown to the right of the images as arbitrary units (log a.u.). Horizontal dashed lines in boxplots indicate background signal for secondary antibody. (***) p<0.001. (C and D) Gene set enrichment analysis (GSEA) results using R5020- and E2-treated Ishikawa expression profiles as discrete phenotypes for classification of normal endometrium (proliferative and secretory) samples. Enrichment profile (green) shows correlation of normal samples at the top or bottom of a ranked gene list (phenotypes). Normalized enrichment scores (NES) and nominal p values (nom.p) are shown in the graphs. (E) Ishikawa cells transfected with an MMTV-Luciferase reporter gene and treated with vehicle (OH) and R5020 10nM (R5020) for 18h. Diagram at the top depicts MMTV LTR promoter features, including several hormone response elements (HRE) and a nuclear factor 1 (NF1) binding site within nucleosome B (dark grey circle and magnification). Numbers in the diagram indicate base pair position relative to transcription start site (TSS). Results are expressed as relative units (r.u.) of Luciferase activity. (F) Representation of *EGFR* TSS and the enhancer sequence located 50kb upstream used to evaluate PR recruitment. Black arrows indicate position of qPCR primers employed on samples treated or not (0) with R5020 for 5, 30 and 60min. Unspecific immunoprecipitation of chromatin was performed in parallel with normal rabbit IgG (IgG). Results are expressed as %input DNA and bars represent mean fold change in PR enrichment relative to time 0 (untreated cells) ± SE of two independent experiments. (*) p<0.05.

Ishikawa cells contain isoforms A and B of PR (PRA and PRB), both of which increased their steady state levels by treating cells with E2 10nM for 12h (Supplementary Fig. S1E and S1F). Pretreating cells with E2 for 12h (preE2) had little effect on the proliferative response to R5020 (Figure 1A), while E2 pretreatment for 48h significantly increased the proliferative effect of E2 exposure, compared to non-pretreated cells (FC 1.47±0.08 v. no preE2). The percentage of cells exhibiting nuclear localization of PR increased upon E2 pretreatment prior to R5020 exposure (T0). Upon exposure to R5020 for 60min the percentage of cells exhibiting nuclear PR was not affected by E2 pretreatment, though the intensity of the fluorescence signal increased in E2-pretreated cells (Figure 1B). Ishikawa cells express considerably higher levels of ERalpha than of ERβ (Supplementary Fig. S1G), suggesting that the proliferative effect of E2 was mediated by ERalpha. R5020 increased nuclear ERal-pha, suggesting a functional PR-ER crosstalk in response to hormonal stimuli (Supplementary Fig. S1H). Such interactions have already been proven in breast cancer T47D cells (Ballaré et al., 2003) and in UIII rat endometrial stromal cells (Vallejo et al., 2005), though in the latter PR remains strictly cytoplasmatic.

Treatment with hormones during 12h produced transcriptomic changes consistent with the physiological stages of normal cycling endometrial tissue (Chi et al., 2020). RNAseq results from Ishikawa cells exposed to E2 10nM for 12h showed a significant resemblance to proliferative endometrium (Figure 1C), while 12h treatments with R5020 10nM regulated a gene expression profile similar to a midi secretory phase (Figure 1D). In line with these i findings, among the top overrepresented bioi logical processes for E2-treated Ishikawa cells i showed angiogenesis and positive regulation of smooth muscle cell proliferation and for R5020-treated cells processes like protein targeting to Golgi and SRP-dependent cotranslational protein targeting to membrane were found (Supplemenatry Fig. S2A). In addition, the majority of regulated genes (81% of R5020 and 63% of E2) were not shared by both hormones (Supplementary Fig. S2B). Genes like *PGR* (progesterone receptor) and cell-cycle regulator *CCND2* (cyclin d2) were upregulated by E2 but not by R5020, while TGFA (transforming growth factor alfa) was upregulated by both hormones (Supplementary Fig. 2B and C).

### Binding of PR and ERalpha to the Ishikawa endometrial cancer genome

To explore the genome-wide distribution of PR and ERalpha binding (PRbs and ERbs respectively) in Ishikawa cells, ChIPseq was performed in different conditions (Figure 2A and Supplementary Fig. S4). First, we analyzed untreated cells (T0) and cells exposed for 5, 30, and 60min to 10 nM R5020 using a specific antibody to PR that detects both isoforms PRA and PRB. Results showed robust PR binding after 30 min of R5020 treatment (R5020 30min) with 1,446 sites, of which 322 sites (22%) were present in untreated cells (PRbs at time zero, T0=331). After 60min of treatment with R5020 (R5020 60min), the majority of sites identified at 30min were still evident (78%), with 336 sites gained and 307 sites lost (Figure 2A and Supplementary Fig. S4A). The representation of PREs in 22% of the PR binding sites that were lost between 30 and 60 min of R5020 treatment was analyzed taking into account common, and unique 30min or 60min PRbs. De novo motif discovery, analysis of information content and quantification occurrences of PRE motifs in such regions did not show differences in the information content (the strength of PRE motif), nor new motif different from PRE, but revealed a higher abundance of PREs in common and unique 60min datasets, yielding 1.72 fold and 1.78 relative unique sites in 30min respectively. Thus PR could bind as monomer isoforms at 30min and as dimer isoforms at 60min, providing more probability of active PR at 60 than at 30min of R5020 treatment. qPCR performed on six regions in the vecinity of hormone regulated genes and occupied by PR at 30 and 60min of R5020 exposure validated ChIPseq results (Supplementary Fig. S4B). These regions were selected according to differentially expressed genes from RNAseq data and top-ranked by peak signal. These results indicate that hormone-dependent PR occupancy increased 5-fold by 30min and stabilized between 30 and 60min of treatment, in accordance with qPCR results (Supplementary Fig. S4C).

**Figure 2.**
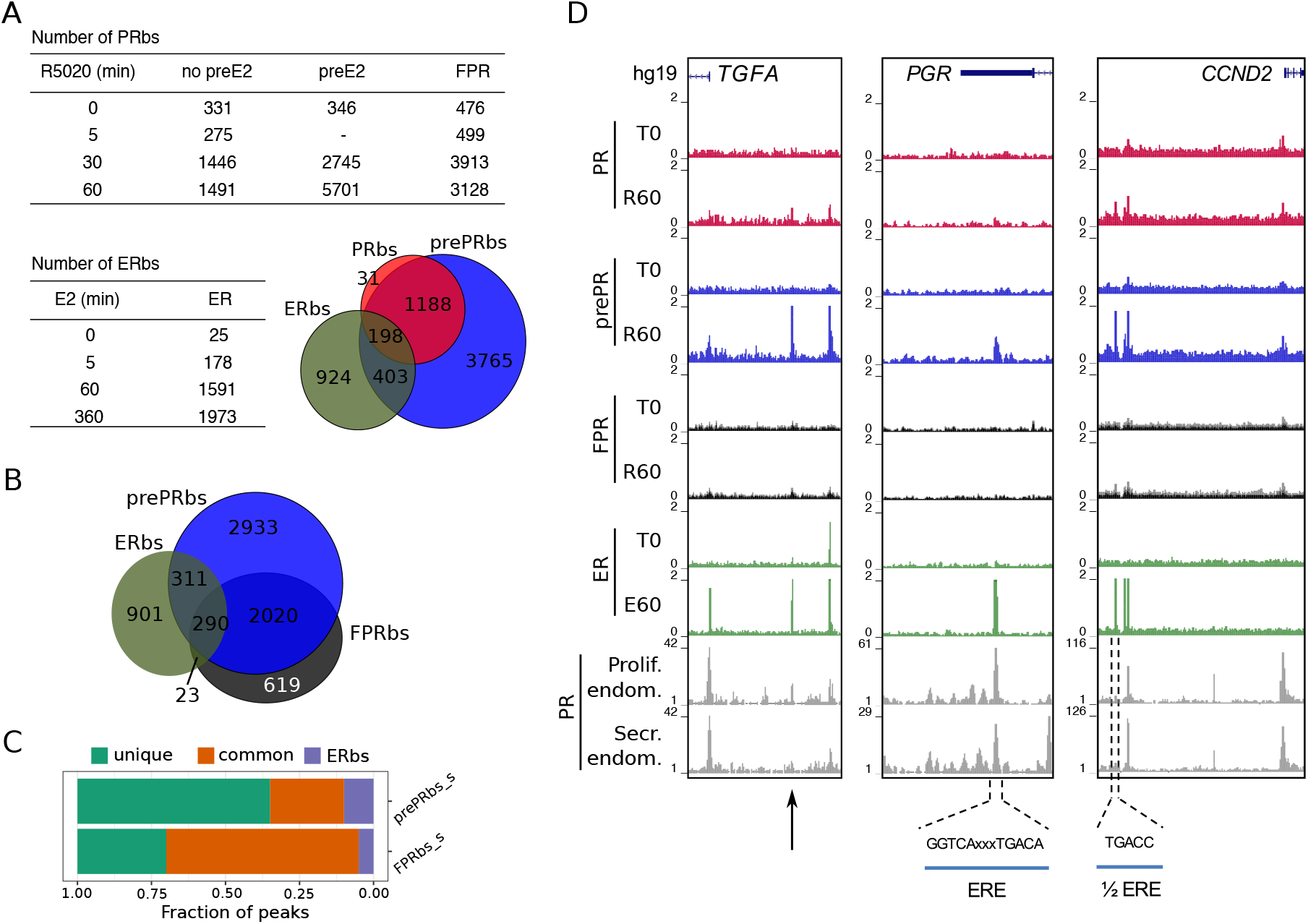
Estradiol induces R5020-dependent PR binding to specific regions in chromatin. (A) Upper table shows total number of PRbs obtained by ChIPseq for untreated (0min) and R5020-treated (5, 30 and 60min) endometrial Ishikawa cells under three different conditions: non-pretreated with E2 (PR), pretreated with E2 for 12h (prePR) and exogenous expression of PR (FPR). Lower table shows number of ERbs using anti-ERalpha antibody on untreated (0min) and E2-treated (5, 60 and 360min) Ishikawa cells. Venn Diagram shows shared binding sites among PRbs (red), prePRbs (blue) and ERbs (green) at 60min. (B) Venn Diagram shows intersection between ERbs (green), FPRbs (dark grey) and prePRbs (blue) at 60min. (C) Fraction of peaks in FPR and prePR after substraction of shared PRbs (FPR_s and prePR_s, respectively) that are not shared with each other (unique), that are common to each other (common) and that are common with ER (ERbs). (D) Normalized coverage of PR and ERalpha binding in untreated (T0) and 60min hormone-treated (R60 and E60) Ishikawa cells and PR binding in proliferative (GSE1327133) and secretory (GSE1327134) endometrium. Black arrow indicates peak of interest. R60: 60min R5020 10nM; E60: 60min E2 10nM. The three regions displayed include *TGFA, PGR* and *CCND2* genes (indicated at the top). An estrogen response element (ERE) and a half ERE are indicated below the peaks.

Next, we explored the recruitment of ERalpha to chromatin of Ishikawa cells exposed to E2 (10nM) for 5, 60 and 360min. Poor ERalpha binding was detected at T0 (25 sites), of which 90% remained occupied throughout all times of treatment with E2. Exposure to E2 resulted in the detection of 178 ERalpha binding sites (ERbs) at 5min, 1,591 at 60min and 1,973 at 360min (Figure 2A and Supplementary Fig. S4D). The majority (85%) of ERbs found at 60min was also identified at 360min (Supplementary Fig. S4D). ERalpha binding at 0, 60 and 360min of E2 treatment was confirmed by qPCR on four of the sites identified (Supplementary Fig. S4E). ChIPseq results point to a clear and sustained E2-dependent enhancement of ERalpha binding (Supplementary Fig. S4F).

De novo motif discovery confirmed that PR binding occurred mostly through PREs exhibiting the complete palindromic response elements (Supplementary Fig. S4G), while ER binding sites were enriched in half-palindromic ERE motifs (Supplementary Fig. S4H). Comparison with previous findings in T47D cells (Nacht et al., 2016) enabled clustering of both PRbs and ERbs into two clases (Supplementary Fig. S4G and S4H, respectively): sites specific for Ishikawa cells (group I; 595 PRbs, group III: 1101 ERbs) and sites present in both Ishikawa and T47D cell lines (group II: 896 PRbs; group IV: 490 ERbs). Classification revealed that PR binds through complete PREs regardless of cell line identity, but in Ishikawa cells ERalpha binds mostly sites with only half of the characteristic palindrome.

### Estrogenic environment defines the landscape for PR binding to the endometrial genome

Shifts in the synthesis and secretion of the ovarian steroids (estrogen and progesterone) during the menstrual cycle serve as the principal hormonal drivers for endometrial changes. Rising circulating estradiol during the mid-to-late follicular phase of the cycle promotes the proliferation of the functional endometrium, and higher E2 levels upregulate *PGR* gene expression (Graham et al., 1995; Kraus and Katzenellenbogen, 1993). A similar result was reported in Ishikawa cells treated with E2 (Diep et al., 2016). To explore the effect of E2 on PR binding to DNA we performed PR ChIPseq analyses on Ishikawa cells exposed to E2 10nM for 12h (preE2) before treatment with R5020 for 30 and 60min. Pretreatment with E2 significantly increased the number of R5020-dependent PRbs (prePRbs), which included most of PRbs already identified in non-pretreated Ishikawa cells (Figure 2A, Table and Venn Diagram). Quantitative real-time PCR validations performed on 6 sites occupied by PR confirmed positioning of the receptor in both non-preE2 (non E2 pretreatment) and preE2 conditions (Supplementary Fig. S5A). It also showed that E2 pretreatment augments both number of PRbs and occupancy of the receptor (signal). Contrary to PRbs in non-pretreated cells, the number of PRbs doubled between 30 and 60min of R5020 in preE2 cells, reaching 5,701 sites (Figure 2A and Supplementary Fig. S5B, S5C and S5D).

Sequencing experiments performed on T47D cells exposed to 10nM R5020 revealed over 25,000 PRbs (Ballaré et al., 2013; Nacht et al., 2016), likely reflecting the high content of PR in these cells. However, a large proportion of these PRbs was considered functionally irrelevant as indicated by the lack of nucleosome remodelling in response to hormone treatment (Ballaré et al., 2013). More recent experiments in T47D exposed to subnanomolar R5020 revealed that around 2,000 PRbs are sufficient to evoke a functional response (Zaurin et al, 2020, personal communication). Hence, the number of PRbs found in Ishikawa cells probably reflects the low concentration of PR, which is compatible with a functional response to progestins. To test this possibility we increased the levels of PR in Ishikawa cells by expressing a recombinant FLAG-PR vector (Supplementary Fig. S5E). These cells, FPR Ishikawa (FPR), expressed levels of PR comparable to T47D cells (Supplementary Fig. S5F) and showed no impairment in hallmark phosphorylation of serine 294 in PR (Supplementary Fig. S5G), indicating that FPR cells were capable of responding to hormone. Upon hormone exposure, FPR cells exhibited rapid binding of PR to the *EGFR* enhancer sequence (Supplementary Fig. S5H). ChIPseq experiments after R5020 exposure showed twice the number of PRbs in FPR cells compared to parental Ishikawa cells. The majority of PRbs identified in Ishikawa cells (>90%) were also detected in FPR cells (Supplementary Fig. S5I), meaning that PR overexpression reflected mostly on an increase in number of binding sites.

Upon hormone induction, sites engaged by PR in Ishikawa cells were also occupied in FPR and pretreated cells, denoting a strong similarity between them (Supplementary Fig. S5I). Although a small number of binding sites was shared between ERalpha and PR in all three conditions, PR binding in pretreated cells exhibited a higher degree of similarity to ER-alpha binding than FPRbs (Figure 2B). Moreover, subtracting PRbs from FPRbs (FPR_s) and prePRbs (prePR_s) heightens this difference, with a much larger fraction of binding sites shared with ERalpha in the case of prePRbs (Figure 2C). Among these sites, one located close to the promoter of *TGFA* gene, identified as an ERbs, showed significant PR binding only in preE2 Ishikawa cells, but not in FPR (Figure 2D, left panel). ERE-containing ERbs, such as the ones found in the transcription termination site of *PGR* gene and immediately upstream of *CCND2* promoter, were occupied by R5020-bound PR in preE2 Ishikawa cells (Figure 2D, middle and right panels). These three genes were upregulated by E2 treatment in RNAseq experiments performed on Ishikawa cells, while only *TGFA* was also upregulated by R5020 (Supplementary Fig. S2C).

The distribution of PRbs and ERbs in non-pretreated Ishikwa cells, in FPR cells and cells pretreated with E2 (prePRbs) relative to TSS of regulated genes was consistent with previous reports in oher cell lines (Ballaré et al., 2013; Need et al., 2015), in that they were enriched in intronic and distal intergenic regions (Figure 3A). Nearly 50% of binding sites localized to distal regions (>50Kb) and approximately 30% to introns other than the first intron, indicating that regulation of gene expression by the steroid receptors PR and ER-alpha is not mediated through proximal promoters but mostly by distal enhancer/silencer sequences. We corroborated these results employing another strategy based on binding site-gene association using the GREAT web tool (see Methods for further details (McLean et al., 2010)). First, we defined a set of genes associated to binding sites with a basal plus extension rule (extended up to 100kb away) and then we intersected this group of genes with R5020- or E2-regulated genes. Of the 1,886 genes regulated by R5020, only 224 (12%) were potentially associated to PRbs, while only 199 of the 950 genes regulated by E2 (21%) proved to be associated to ERbs (Figure 3B).

**Figure 3.**
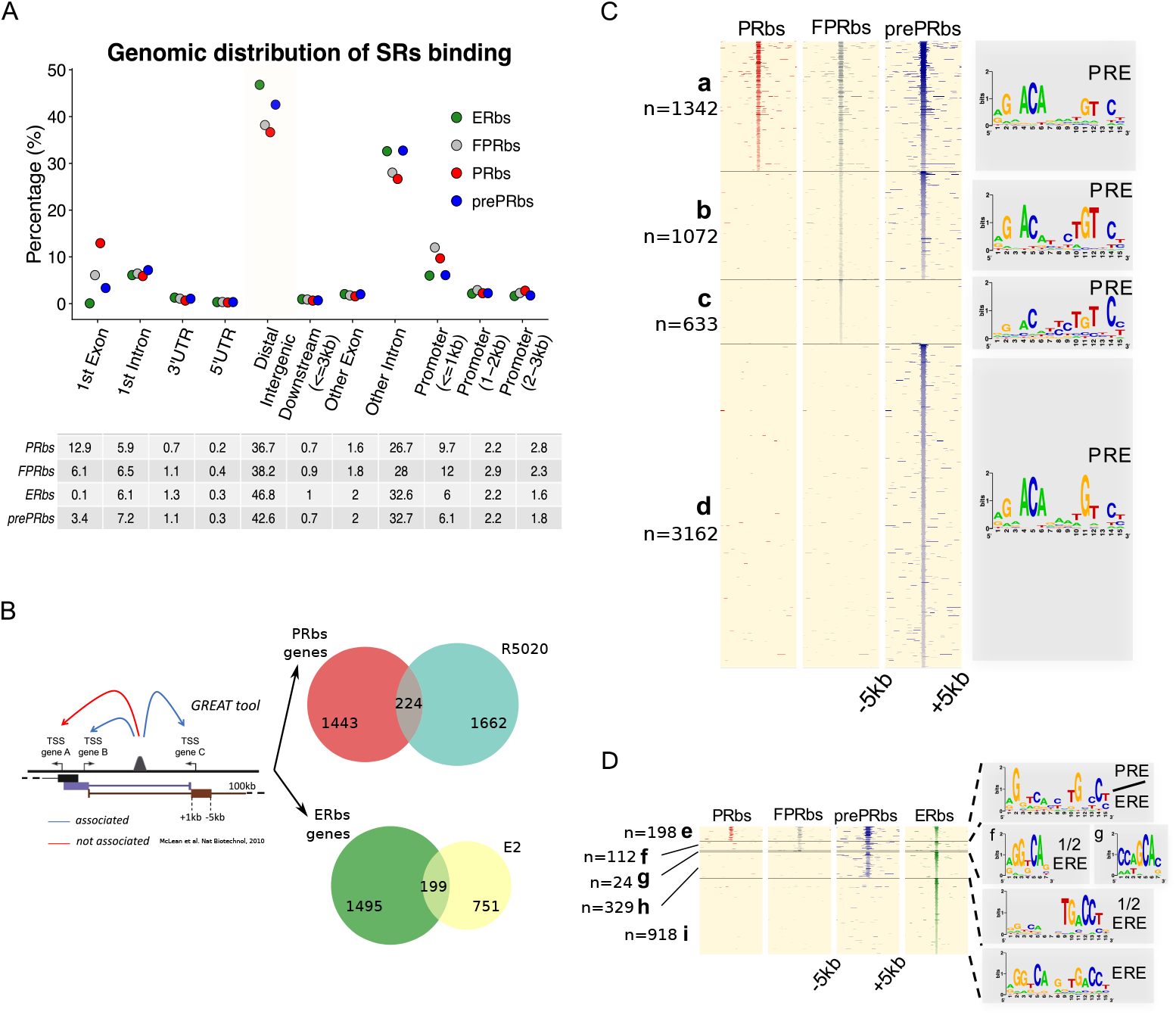
A fraction of E2-induced PRbs localize on ERbs and contain half ERE motifs. (A) Classification of steroid receptor binding relative to genomic features expressed as percentage (%) of peaks after 60min of hormone treatment inside each feature. Legend at the top right corner indicates the color key for ERbs (green dots) and three conditions of PR binding: non-pretreated with E2 (PRbs, red dots), pretreated with E2 for 12h (prePRbs, blue dots) and exogenous expression of PR (FPRbs, grey dots). The table below shows percentages represented in the plot. (B) To the left: Representation of GREAT tool association rules adapted with modifications. To the right: Venn diagrams show intersection between PRbs-associated genes and R5020-regulated genes (top), and ERbs-associated genes and E2-regulated genes (bottom). (C) Peak signals in PRbs, FPRbs and prePRbs from 60min R5020-treated Ishikawa cells were plotted as heatmaps. Regions were defined inside a window of 10kb centered in peak summit (±5kb) and intensity of the signal correspond to number of reads in each region. Heatmap is subdivided into 4 mutually exclusive groups depending on shared/partly shared/non-shared binding sites: a (n= 1342), sites shared by all three conditions of PR binding; b (n=1072), sites uniquely found in FPR and prePR; c (n=633), sites found only in FPR; and d (n=3162), sites found only in prePR. De novo motif discovery (MEME) was performed on all groups and results are indicated as sequence logos to the right of the map, including the name of the most related known motif. PRE: progesterone response element. (D) Peak signals in PRbs, FPRbs and prePRbs as in (C), and ERbs from 60min E2-treated Ishikawa cells. Heatmap was subdivided into 5 mutually exclusive groups: e (n=198), sites shared by all three conditions of PR binding and ER binding; f (n=112), sites shared by FPRbs, prePRbs and ERbs; g (n=24), sites shared by FPRbs and ERbs; h (n=329), sites shared by prePRbs and ERbs; and i (n=918), sites uniquely found in ERbs. Motif discovery was performed as in A for all groups and results are shown to the right of the map, including the most related known motif. ERE: estrogen response element; 1 /2 ERE: half ERE.

As expected, from the sequences contained in 10kb windows centered in peak summits of PRbs, FPRbs and prePRbs, the PRE emerged as the most representative binding motif (Figure 3C), including sites uniquely found in FPR (group c: 633) or preE2 (group d: 3,162) cells. While comparison between ERalpha and PR ChIPseq results showed few similarities regarding identity of binding sites, with a set of 216 shared by both hormone receptors, pretreatment with E2 added nearly twice as many binding sites to the pool shared with ERalpha (from Figure 2A, Venn Diagram). The most representative motif discovered in these sites - only shared by ERalpha and prePR-was a half ERE (Figure 3D, group h: 329) that was highly similar to the motif observed in sites uniquely found in Ishikawa ERbs (from Supplementary Fig. S4H, group III). Sites shared by ERalpha and PR in all three conditions resulted in an unclear combination of PRE and ERE motifs (Figure 3D, group e-g). Degenerated motif logo in group g showed no association to any known motif, probably due to a corrupt analysis performed on insufficient data, and the partially degenerated motif logo in group e showed limited association to both PRE and ERE (PRE/ERE).

Taken together, this evidence suggests that, provided there is an estrogenic background, activated PR could regulate estrogen-dependent Ishikawa-specific transcriptome by binding sites already or formerly bound by ERalpha.

### PAX2 binds chromatin in close proximity to ERalpha and PR binding sites in Ishikawa cells

Evidence described so far partially explains cell type specific hormone-dependent gene regulation, though it is not sufficient to understand the mechanisms underlying differential binding of hormone receptors to chromatin. Initially, we addressed this by contrasting the sequences of ERbs and PRbs from groups IIV, i.e. hormone regulated Ishikawa specific, (from Supplementary Fig. S4G and S4H) with an array of 1,395 known TF binding motifs (see Methods). Results revealed an enrichment (p-value<1e^-4^) of multiple members of the PAX family −including variants 2, 5, 6 and 9-in groups I and III, i.e. In Ishikawa specific PRbs and ERbs (Supplementary Fig. S6A and S6B, respectively), suggesting that members of the Pax family may be involved in PR and ERalpha action in Ishikawa cells. Unbiased comparison (all sites) of enrichment in TF binding motifs between Ishikawa and T47D PRbs showed similar results for PRbs, although the enrichment was less significant (Supplementary Fig. S6C). Moreover, while enrichment of PAX motifs was also observed around ERbs in Ishikawa cells (Supplementary Fig. S6D), this was not the case with T47D cells, in which examples like the well-known breast-related pioneer transcription factor FOXA1, were found instead (Supplementary Fig. S6E).

Enrichment of NR3C1-4 (mineralocorticoid, glucocorticoid, progesterone and androgen receptors) and ESR1 motifs included into the 1,395 known motifs corroborated de novo discovery performed with MEME in both Ishikawa and T47D cells. Stronger enrichment of PAX motifs was observed in prePRbs compared to PRbs (Figure 4A), indicating that PR binding to regions potentially bound by PAX is favored after E2 pretreatment. Coherently, while equivalent fold enrichment values were detected when comparing prePRbs to ERbs (Figure 4B), comparison between prePRbs and FPRbs showed that increased PR levels alone were not sufficient for a greater association to PAX binding motifs (Supplementary Fig. S6F). Consistently, RNAseq experiments on Ishikawa cells treated either with R5020 10nM or E2 10nM for 12h showed putative PAX2 binding sites among the top 20 significantly enriched TFs (DAVID web-based tool (Huang et al., 2009)) on differentially regulated genes (Figure 4C). ER was also predicted to bind on E2-responsive genes, while glucocorticoid receptor (GR) motif (PR-like motif) was detected on R5020-responsive genes.

**Figure 4.**
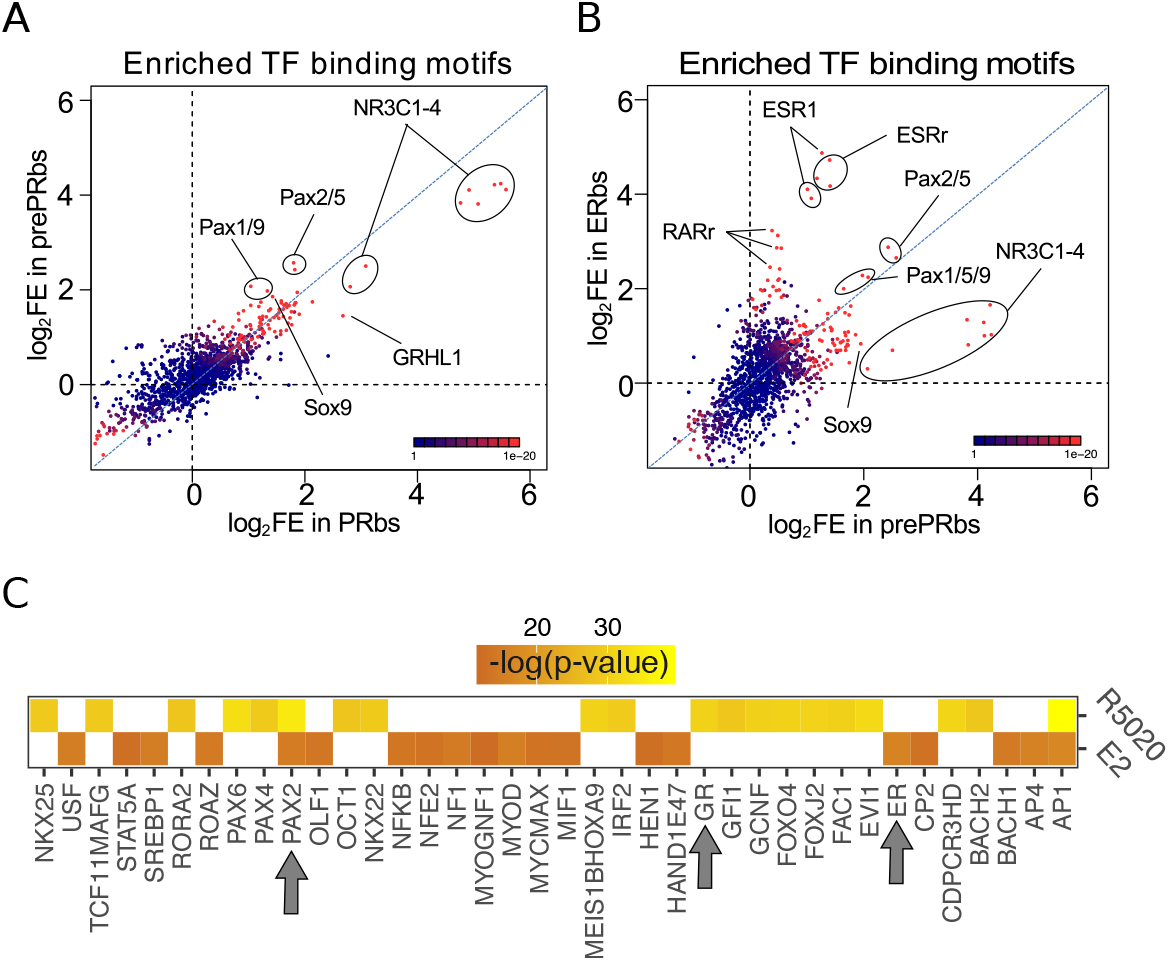
Putative PAX2 binding sites are associated with PR and ERalpha binding and hormone-regulated genes in Ishikawa cells. (A) Fold enrichment values (log2FE) of 1.395 known TF binding motifs on prePRbs and PRbs. Combined p-values for enrichment analyses are indicated through the color key displayed at the lower right corner of the plot. Relevant motifs pointed on the plot correspond to NR3C1-4, members of the PAX family (1, 2, 5 and 9) and SOX9. (B) Comparison as in (A) between prePRbs and ERbs. Relevant motifs pointed on the plot correspond to NR3C1-4, members of the PAX family (1, 2, 5 and 9), SOX9, ESR1 and estrogen related (ESRr) and retinoic acid receptor (RARr). (C) Predicted UCSC Transcription Factor (TFBS) binding on genes regulated by 12h treatments with R5020 10nM and E2 10nM in Ishikawa cells were analysed using DAVID web-based functional enrichment tool. Heatmap shows the top 20 TFBS predicted (p<0.05)for R5020- and E2-regulated genes from RNAseq results expressed as −log(p-value). Arrows indicate position of PAX2, GR (PR-like binding motif) and ER.

PAX association to PR and ERalpha action was also evaluated by immunofluorescence against PAX2. Nuclear localization of PAX2 was observed predominantly after 60min of R5020 in pretreated and non-pretreated PR+ cells (Figure 5A), indicating that hormonal treatment promotes co-localization of PAX2 and PR in nuclei of Ishikawa cells. Similar results in PAX2 localization were obtained after treating Ishikawa cells with E2 for 60min (Supplementary Fig. S6G). The increase in nuclear PAX2 signal is not due to changes in protein levels, which were not affected by treatment with either R5020 or E2 (Supplementary Fig. S6H). In accordance to motif analysis results, PAX2 was not detected in nuclei of T47D cells after hormonal treatments (Supplementary Fig. S6I).

**Figure 5.**
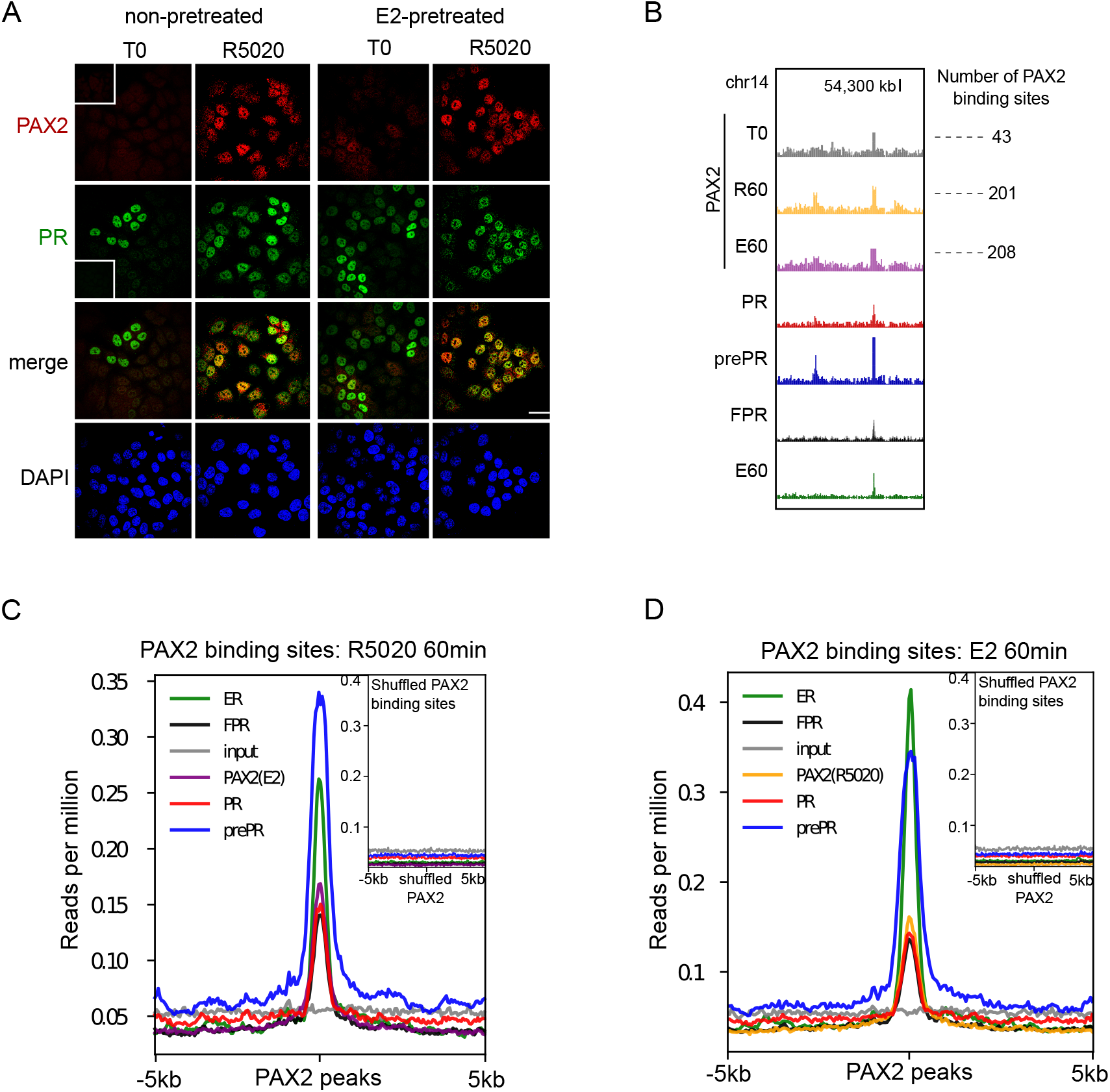
PAX2 co-localizes with PR and ERalpha in nuclei of Ishikawa cells and it is positioned primarily in the vicinity of receptors binding sites. (A) Immunofluorescent detection of PR (green) and PAX2 (red) in untreated (T0) and 60min R5020-treated (R5020) Ishikawa cells which were pretreated or not with E2 for 12h (non-pretreated, E2-pretreated). Images were merged for co-localization analysis (merge). Scale bar is shown in the panels and is equivalent to 30*μ*m. (B) PAX2 binding profile and peak calling output (thicks below peaks) inside a region of 70kb of chromosome 12. Number of PAX2 binding sites for untreated Ishikawa cells and treated with R5020 for 60min or E2 for 60min is shown to the right of the profiles. Tracks for PRbs, prePRbs, FPRbs and ERbs are displayed below the profiles for the same region. (C) Binding profiles of ER (green), PR (red), FPR (black) and prePR (blue) on PAX2 binding sites of 60min R5020-treated Ishikawa cells. PAX2 binding after 60min E2 treatment was included (purple). Inset shows signal profiles centered on shuffled R5020-dependent PAX2 binding sites. (D) Binding profiles as in (C) on PAX2 binding sites of 60min E2-treated Ishikawa cells. PAX2 binding after 60min R5020 treatment was included (orange). As in (C), inset shows signal profiles centered on shuffled E2-dependent PAX2 binding sites.

To extend these findings, we performed PAX2 ChIPseq experiments on untreated cells and in cells exposed for 60min to either R5020 or E2. The results confirmed PAX2 binding to chromatin following hormonal treatment (Figure 5B). Even though identified PAXbs were few (T0: 43, R60: 201 and E60: 208), most of PAX2 binding occurred after R5020 and E2 treatments. Moreover, PAX2 binding was not stochastically distributed in the genome of Ishikawa cells but rather partially associated to ERbs and PRbs. This association was stronger for PR binding in cells pre-teated with E2 than in non-pretreated cells or in cells overexpressing recombinant PR (Figure 5C). Similar results were observed for ERbs in response to E2 (Figure 5D), indicating that PAX2, and possibly other members of the PAX family may co-operate with PR and ERalpha for binding to chromatin in Ishikawa cells but not in T47D cells, in which neither enrichment for PAX binding motif nor nuclear localization of PAX2 was detected.

### Under estrogenic conditions, PR and PAX2 conform endometrial regulatory domains in open chromatin compartments

Nuclear architecture is a major determinant of hormonal gene regulatory patterns (Le Dily et al., 2014). Therefore, we used in nucleo Hi-C technology to study the folding of chromatin across the genome of Ishikawa cells by generating genome-wide contact datasets of cells untreated (T0) or pretreated with E2 for 12h, and exposed to R5020 or E2 for 60min. A comparison of contact matrices at 20 kb resolution of untreated Ishikawa cells to T47D cells confirmed the high degree of conservation on the borders of topologically associating domains (TADs) (Supplementary Fig. S7A). TADs are grouped into two chromatin compartments A and B, which represent the active open chromatin (A) and the closed inactive chromatin (B) respectively. Analysis of such compartments showed a cell type-specific patterning (Supplementary Fig. S7B), in which Ishikawa samples from two independent experiments were more closely related to each other than any of them to a T47D sample (Supplementary Fig. S7C and S7D). However, A/B profile distribution in Ishikawa cells was independent from hormonal treatments (Supplementary Fig. S7B and S7E), meaning that chromatin was in a primed state that conditioned hormone-dependent regulation of gene expression. Detailed analysis revealed that 7% of A domains in Ishikawa cells were B in T47D cells, and 12% of B domains in Ishikawa cells were A in T47D cells (Supplementary Fig. S7F). A total of 861 genes encompassed in the A compartment in Ishikawa cells belong in the B compartment in T47D cells, and 1,438 genes in B compartaments in Ishikawa cells belong in A in T47D cells (12%), suggesting that distribution of A and B compartments could in part explain cell type specific gene expression profiles.

To evaluate whether chromatin states are related to gene expression through differential binding of hormone receptors to DNA, we intersected PR and ERalpha ChIPseq results with the A/B compartment coordinates. Both transcription factors, PR and ERalpha, bound A compartments more frequently than B, meaning that open genomic regions in Ishikawa showed preferential binding of the hormone receptors (Supplementary Fig. S7G). Neither pre-treatment with E2 nor expression of recombinant PR modified the preferential binding of the PR to the A compartments.

As mentioned above, PAX2 binding occurs mostly in close proximity to PR and ERalpha binding sites. In fact, distances between PAXbs and PRbs were remarkably shorter in E2 pretreated cells than in any other condition (Supplementary Fig. S7H). This raised the question of whether recruitment of PR together with PAX2 to open chromatin compartments facilitates regulation of gene expression. To study this notion, we defined putative endometrial regulatory domains that we named “Progestin Control Regions” (PgCR) with the capacity to potentially regulate nearby genes. The restrictions for being a regulatory domain, which consisted in containing at least two PRbs separated by a maximun distance of 25kb and a PAXbs (represented in Figure 6A: PgCRs Definition), were met mostly under E2 pretreated conditions. This outcome was due to the strong association between prePRbs and PAXbs, though it may have been aided by the increased PR protein levels. However, the sole increment in PR protein levels was not enough to force an association to PAXbs, given that FPR cells did not show similar results (Figure 6A: PgCRs Definition).

**Figure 6.**
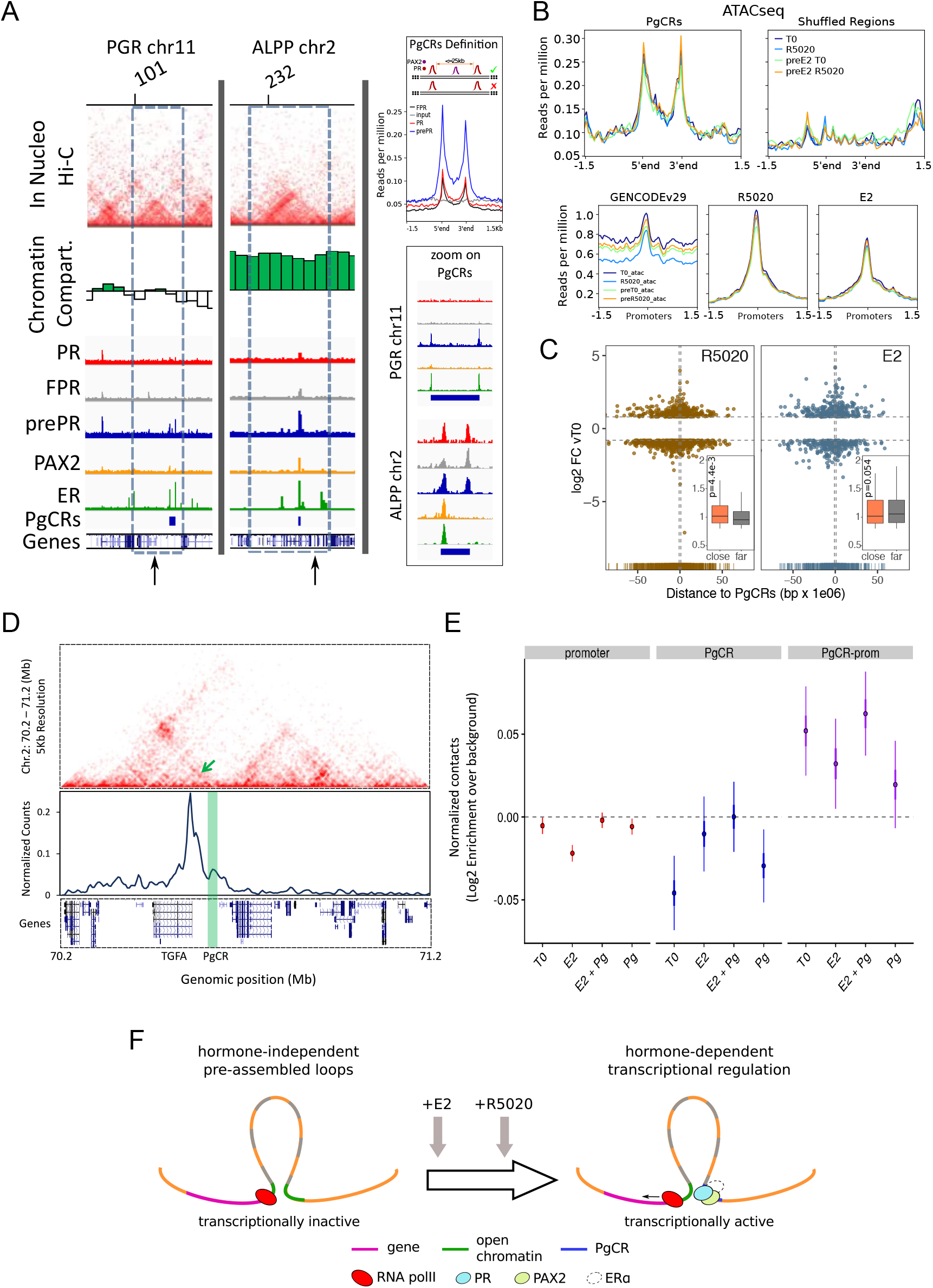
Convergence of PR and PAX2 binding in TADs with regulated genes defines potential endometrial regulatory domains. Convergence of PR and PAX2 binding in TADs with regulated genes defines potential endometrial regulatory domains. (A) Upper panel shows the contact matrices at a resolution of 20kb obtained by In Nucleo Hi-C in *PGR* and *ALPP* loci. Middle panel shows the spatial segregation of chromatin as open or closed compartments inside TADs (green bars: A compartment; white bars: B compartment - see methods section). The bottom panels show ChIPseq signal distribution of PR, FPR, prePR, PAX2 and ERalpha as well as the location of PgCRs and genes over the region. The dashed rectangle restricts the TAD of interest and the vertical arrow marks the TSS of *PGR* and *ALPP*. Definition of PgCR: Coverage profiles of PR (red), FPR (black) and prePR (blue) binding on Progesterone Control Regions (PgCRs) delimited by the start and end labels, and flanked upstream and downstream by 1.5Kb regions. Input sample (grey) was included in the plot. Rules for qualifying as a control region are depicted on top of the profile plot. Magnified images over Control Regions are shown to the right (zoom on PgCRs). (B) ATACseq peaks from cells untreated (T0), treated with R5020 for 60min, 12h E2-pretreated (preE2 T0) and E2-pretreated followed by 60min treatment with R5020. Signal was plotted over Control Regions, shuffled Control Regions (Shuffled Regions), promoters of all annotated genes from GENCODE database (GENCODEv29) and promoters of genes regulated by 12h treatments with R5020 or E2. (C) Plot shows fold change values of genes regulated by R5020 and E2 (v. untreated cells) relative to Control Regions. Genes located upstream of PgCRs are represented with negative distance values. Dashed horizontal lines mark fold change cut-off points (|log2FC|=0.8) and vertical lines are placed at position −1 and 1Mb. Insets depict comparison of fold change values (absolute values) between genes located beneath (close) and over (far) a 1Mb distance from PgCRs. Statistical significance for this comparison was determined with Welch Two Sample t-test and is represented by a p value on the plot. (D) Top panel: Hi-C contact map at 5kb resolution of Chromosome 2 (70,200,000-71,200,000) obtained in Ishikawa cells and showing the organization around TGFA gene locus. Middle panel: Virtual 4C profile at 5kb resolution (expressed as normalized counts per thousands within the region depicted above) using the TGFA promoter as bait and showing the contacts engaged between TGFA promoter and the PgCR detected in this region (highlighted in green). Arrow on top panel highlights the position of the loop in the map. Bottom panel shows the positions of genes in the region depicted. (E) Distributions of observed versus expected interactions established between promoters (red - left), between PgCRs (blue - middle) and between Promoters and PgCRs (purple - right) located within a same TAD in Ishikawa cells treated as indicated below. (F) Representation of a chromatin loop involving a PgCR and the promoter of a regulated gene. Initially, the gene is transcriptionally inactive even though the loop is already formed. After hormone induction (E2 pretreatment followed by R5020), PR, PAX2 and in some cases ERalpha occupy open chromatin compartments in contact with promoters resulting in transcriptional activation.

Considering that TAD borders may act as regulatory barriers, we removed from further analysis any region that, in spite of satisfying the rules for being a PgCR, was localized across a barrier as well. In agreement with this restriction, the sizes of PgCR −with an average of 25kb-were smaller than TADs −with an average of 1000kb- (Supplementary Fig. S7I). In addition, the majority of the 121 identified PgCRs (coordinates in hg38 can be found as Supplementary Data) were not located near the TAD borders, but in the TAD center (Supplementary Fig. S7J), where most non-housekeeping genes are found (Le Dily et al., 2019). Moreover, PgCRs seem to be located in A compartments in the vecinity of hormone-regulated genes like *PGR* and *ALPP* (Figure 6A). Expression of these genes was analyzed by qPCR of total RNA samples of Ishikawa cells exposed to hormone for 12h, which showed that *ALPP* is induced by both hormones and *PGR* is only induced by E2 (Supplementary Fig. S7K).

As was mentioned before, the Hi-C matrices were used to determine the spatial segregation of chromatin in both open and closed chromatin compartments (A/B), and the A:B ratio was independent of hormone treatment. Consistent with these results, ATACseq signal on PgCRs remained unchanged upon hormone exposure, but it decreased after shuffling the coordinates for PgCRs, indicating that chromatin was readily and non-randomly accessible to TFs in these locations (Figure 6B, top panels). Although ATACseq peaks were also detected on promoters of hormone-regulated genes, the signal did not differ after hormone exposure (Figure 6B, bottom panels), implying that treatments were not responsible for opening the chromatin in these regions. In addition, both R5020- and E2-regulated genes with highest FC values (v. T0) were concentrated under 1Mb (“close”) away from PgCRs (Figure 6C), though the comparison between FC values of “close” and “far” (over 1Mb) regulated genes was significant only in the case of R5020 (p=4.4e^-3^; Figure 6C, inset).

Further analysis on Hi-C contact matrices revealed that PgCRs preferentially interact with promoters of hormone-regulated genes (Figure 6D). Although PgCR-promoter interactions were non-random and mostly intra-TAD, we found no difference in contact enrichment between treated and untreated cells (Figure 6E). These results are consistent with ATACseq profiles and imply that chromatin would be pre-assembled into regulatory loops −involving PgCRs and promoters-which are transcriptionally inactive until hormone-dependent binding of steroid receptors and PAX2 triggers PolII activation (Figure 6F).

These results suggest that specific binding of PR, PAX2 and ERalpha to chromatin occurs in compartments that are present in a permissive (open) or restrictive (closed) status depending on the cell line, and are not modified by short term hormone exposure (Figure 6F). However, it is not yet clear the role of PAX2 in PR binding to PgCRs. Summing up, PR and ER bind mostly to non-common sites that exhibit the corresponding consensus sequences, and are adjacent to PAX2 binding. Therefore, the endometrial specific hormone response results in part from specific chromatin compartments, unique receptor binding sites and selective TFs binding partners to regulate gene expression.

### Genes contained in TADs with PgCRs are associated to endometrial tumor progression

To explore the possibility that alterations in the expression profile of genes under the influence of PgCRs were related to disease progression such as endometrial cancer, we examined the genes carrying the most frequent mutations in a cohort of 403 cases diagnosed with endometrial adenocarcinoma (data available in The Cancer Genome Atlas, TCGA, Project TCGA-UCEC). The top 1000 most frequent somatic mutations in these cases were distributed among 837 genes, 33 of which belonged to PgCR-containing TADs (Figure 7A), comprising 6% of the 517 protein coding genes that may be regulated by direct interactions with a corresponding PgCR (PgCR-genes). In fact, pathway analysis of these 517 genes revealed a clear bias towards regulation of immunological processes and transcriptional alterations in cancer (Figure 7B), suggesting that PgCR-genes may participate in key steps of tumor onset and progression.

**Figure 7.**
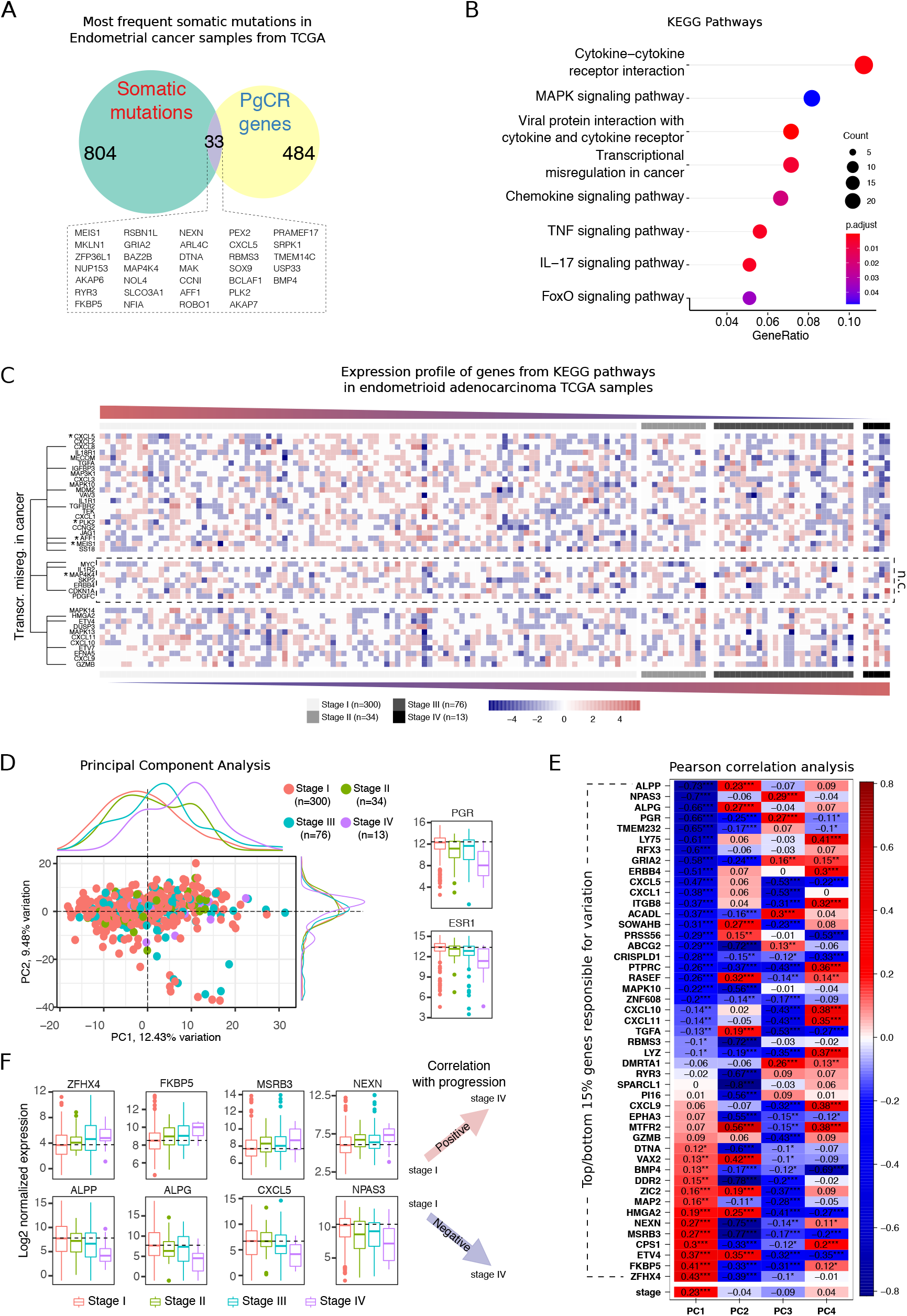
Altered expression of genes contained in TADs with PgCRs correlates with drivers of endometrial tumor progression. (A) Venn diagram of genes carrying the top 1000 most frequent mutations in a cohort of 403 cases of endometrial adenocarcinomas from TCGA (n=837) and protein coding genes contained in TADs with PgCRs (n=517). Names of genes located in the intersection of the two groups (n=33) are detailed below the diagram. (B) Enriched KEGG pathways for all protein coding genes included in TADs containing PgCRs. Number of genes in each category and adjusted p-value (Benjamini-Hochberg) are indicated in the plot. (C) Heatmap of genes from enriched pathways using normalized counts from 423 endometrioid adenocarcinoma samples (TCGA) classified according to the FIGO system (Stage I to IV). Top panels show genes that decrease expression with stage and bottom panels genes that increase expression levels with stage. Genes in the middle panels do not show a clear expression pattern (n.c.: not clear). Each cell in the heatmap represents the mean expression value of three samples (bin). Genes frequently mutated in endometrial adenocarcinomas are marked with an asterisk (*) and genes belonging to pathway transcriptional misregulation in cancer are indicated by a dendrogram. (D) Bi-plot of PCA results depicting scores of components 1 and 2 (PC1 and PC2). Dots represent the samples included in the analysis (n=423) and color identifies the tumor stage. Density marginal plots represent distribution of scores for each stage. (E) Correlation of variables (genes) and stage to principal components (PC1 to PC4). Pearson correlation scores are shown inside the cells and represented in a color scale (red as positively correlated and blue as negatively correlated). The 29 genes displayed in the matrix are included among the 15% of genes that give rise to the variation in principal components (PC1 to PC4). Significant results are indicated in the cells: **** p<0.0001, *** p<0.001, ** p<0.01, * p<0.05. (F) Distribution of normalized counts (log2) for genes positively (top row) and negatively (bottom row) correlated with endometrial cancer stage progression. Dashed line indicates position of the median in Stage I.

We also studied the expression profile of genes involved in enriched pathways (40 genes) using 423 Endometrioid adenocarcinoma RNAseq samples previously classified into stages (Stage I: 300, Stage II: 34, Stage III: 76 and Stage IV: 13) according to the FIGO system (International Federation of Gynecology and Obstetrics). Considering the inherently heterogeneous nature of tissue samples, we intentionally set a permissive fold change cut-off value when comparing Stage I to Stage IV to detect probable subtle differences. The analysis showed that 22 of the 40 genes tended to decrease their expression levels with stage progression (Stage IV v. Stage I: log2FC<-0.5, downregulated genes) including the frequently mutated genes CXCL5, PLK2, AFF1 and MEIS1, 7 genes had no clear tendency and 11 genes increased their expression levels like chemokines CXCL9/10/11 (Stage IV vs. Stage I: log2FC>0.5, upregulated genes) (Figure 7C). Among the genes included in the pathway “Transcriptional misregulation in cancer” (14 genes), there were 7 downregulated genes during tumor progression *-CXCL8, IGFBP3, MDM2, TGFBR2, AFF1, MEIS1* and *SS18*-, 3 genes with ambiguous behavior among samples −MYC, *ILR2* and *CDKN1A*- and 4 upregulated genes in Stage IV tumors −*HMGA2, ETV4, ETV7* and *GZMB*-.

To determine if the PgCR-genes could be drivers of progression in endometrial adenocarcinoma, we performed Principal Component Analysis (PCA) on the 423 Endometrioid adenocarcinoma RNAseq samples using the 517 PgCR-genes as variables. Assessment of PCA results revealed that inter-stage variation was mostly explained within the first two components (PC1: 12.43% and PC2: 9.48%) and notably, this variation was accompanied by a considerable change in PGR mRNA levels (Figure 7D), which could partly account for differences between stages. Although *ESR1* is not directly influenced by PgCRs, we detected that its mRNA levels were also reduced with stage progression. The signature of genes regulated in conjunction with loss of hormonal regulation could assign novel markers in order to differentiate the evolution of malignancies depending on the presence of these molecules to tune a specific response. Finally, we identified the genes that contributed the most to inter-stage variation (top/bottom 15%), confirming that *PGR* (r=-0.66) was indeed negatively correlated with progression (Figure 7E), as well as *ALPP* (r=-0.73), *NPAS3* (r=-0.7), *ALPG/ALPPL2* (r=-0.66) and *CXCL5* (r=-0.47) among others, while *ZFHX4* (r=0.43), *FKBP5* (r=0.41), *MSRB3* (r=0.27) and *NEXN* (r=0.27) were positively correlated with the stage. Correlation results for these genes were consistent with the distribution of their normalized expression values across stages (Figure 7F).

## Discussion

There seems to be consensus that the way in which combinations of TFs assemble their binding sites contributes to the folding of the genome in cell type specific patterns that orchestrate the physiological coordination of gene expression programs required for the proper development and function of complex organisms (Lambert et al., 2018; Stadhouders et al., 2019). There is evidence that the same TF can regulate different gene sets in different cell types (Gertz et al., 2012), but the mechanisms through which hormone receptors regulate endometrial specific gene networks had not been previously deciphered. Here, we describe ERalpha and PR binding to the genome of endometrial cancer cells and analyze their specific chromatin context. In this genomic study we used Ishikawa cells, given that they are a good model of Type I epithelial endometrial cancer [37] containing ERalpha and PR.

It was reported that in Pgr Knockout (PRKO) mice the absence of PR results in unopposed estrogen-induced endometrial hyperplasia (Lydon et al., 1995). As for the two isoforms of PR, the PRB isoform is considered a strong transcriptional activator while PRA can function as a transcriptional inhibitor of PRB activity(Mulac-Jericevic et al., 2000). Selective ablation of PRA in mice results in a PRB dependent gain of function, with enhanced estradiol-induced endometrial proliferation (Conneely et al., 2003). Ishikawa cells express more PRB than PRA, coherent with PRB dominance in glandular epithelial cells (Mote et al., 1999). To explore the mechanism underlying the endometrial specific response to ovarian steroids hormones, we studied the genomic binding of ERalpha and PR by ChIPseq in hormone untreated Ishikawa cells and in cells exposed to hormone for different time periods. We discovered that the majority (67%) of PRbs after estradiol pretreatment were new sites not present in untreated cells and different as well from ERbs occupied after estradiol treatment. Just 639 PR binding sites (11% of all PRbs) were the same for both PR and ERalpha. This indicates that contrary to what was described in breast cancer cells (Mohammed et al., 2015; Singhal et al., 2016), in endometrial cells PR binding has little influence on ERalpha binding. In Ishikawa cells, binding of ER and PR occurs mainly at ERE and PRE sequences, respectively, in regions that are also enriched in PAX response elements. Ishikawa cells are rich in PAX TF and PAX ChIPseq shows a similar overlapping with ERbs and PRbs.

When we analyzed chromatin topology of Ishikawa cells using Hi-C we found that PRbs and ERbs are enriched in Topologically Associating Domains (TADs) containing hormone regulated genes. These TADs were predominantly part of the open (A) chromosome compartment, even in cells not exposed to hormone. This was confirmed by ATACseq results showing that the sites where the hormone receptors will bind were already more accessible for enzyme cleavage, suggesting that hormone independent mechanisms were responsible for the generation and maintenance of the hormone responsive TADs. In that respect, it is interesting that we found an enrichment of PAXbs near PRbs in these TADs containing progesterone regulated genes, suggesting that PAX2 could generate the open chromatin conformation that enables PR binding and facilitates the interacting loops detected in Hi-C experiments. Loss of PAX2 expression has been implicated in the development of endometrial intraepithelial neoplasia (EIN) (Sanderson et al., 2017) and PAX2 is potentially useful in the diagnostic of difficult EIN cases (e.g. where there is no “normal” tissue available to act as an internal control when assessing nuclear morphology) (Quick et al., 2012). Our results connect PR response elements with PAX2 and 3D chromatin conformation, which is consistent with the preservation of progestin regulation in differentiated cancer cells expressing hormone receptors and may be lost in undifferentiated tumor cells, which do not express hormone receptors. We hypothesize that PR-PAX-PR binding sites containing regulatory domains that we name PgCRs could reflect PR shadow enhancers (Cannavo et al., 2016) in endometrial cells.

The redundancy of PRbs associated to endometrial specific gene expression may reinforce a genetic mechanism to ensure progestin regulation in tissue under hormonal influence, in periods in which there is low or no circulating hormone. Notably, the only described super-enhancer in endometrial carcinomas is the Myc super-enhancer and is not hormonally regulated (Zhang et al., 2016). We postulate the existence of a novel subset of 121 strategical endometrial regulatory domains in this hormonally responsive endometrial cancer cell line. Among them the *TGFA* gene presents one of PgCR-promoter interaction that could explain hormone regulation previously reported in this cells (Hata et al., 1993). This concept could be exploited to guide treatments oriented to recover progestin regulation over estrogen proliferative effects in endometrial malignancy.

Previous results in T47D mammary cancer cells have shown Hormone Control Regions, which include ERbs and PRbs acting in conjunction with FOXA1 and C/EBPa (Nacht et al., 2019) interact with promoters of hormone regulated genes in hormone responsive TADs and organize the high level folding of the genome (Le Dily et al., 2019). Although the analysis of interaction between PgCR and different ERalpha enriched binding regions in endometrial cells remains to be performed, our present study proposes that PR binding sites originated under estrogenic conditions and acting in conjunction with PAX2, fulfil a similar function in differentiated hormone-responsive endometrial cancer cells. Thus combinations of the same hormone receptors and different transcription factors account for cell type specific expression of different gene regulatory networks in part by generating and maintaining different genome topologies.

Droog et al. highlights that “the divergence between endometrial tumors that arise in different hormonal conditions and shows that ERalpha enhancer use in human cancer differs in the presence of nonphysiological endocrine stimuli” (Droog et al., 2017). They reported that ERalpha-binding sites in tamoxifen-associated endometrial tumors are different from those in the tumors from nonusers. It has yet to be explored whether the response to progesterone and sinthetic progestins, used in treatments of hormone-dependent endometrial cancers, is affected by the changes resulting from the use of tamoxifen.

On the other hand, estrogen receptor a (ER) and glucocorticoid receptor (GR) are expressed in the uterus and have differential effects on growth (Vahrenkamp et al., 2018). Expression of both receptors was associated with poor outcome in endometrial cancer and the simultaneous induction of ER and GR leads to molecular interplay between the receptors (Vahrenkamp et al., 2018). In our conditions, R5020 induces genes with GR/PR putative binding sites, enabling regulation that could result in a similar ER-GR pathological outcome.

Regarding genes under PgCRs regulation, *ETV4* was one of the most frequent genes encompassed in a PgCR giving rise to the variation of PCA applied to endometrial adenocarcinoma tumors. This gene was recently reported as playing a major role in controlling the activity of ER and the growth of endometrial cancer cells (Rodriguez et al., 2020). Like *ETV4*, other genes such as *MEIS1, ZFHX4, FKPB5, TGFBR2* are under regulation of PR specific endometrial enhancers present in PgCRs and could be responsible for the advance in malignancy of endometrial cancer through a progressive repression of the immune response together with an increased EMT-based metastatic/invasive potential (Bhanvadia et al., 2018; Alfaro et al., 2017; Bai et al., 2019; Cancer Genome Atlas Research Network et al., 2017; Monsivais et al., 2019; Dufait et al., 2019; Ma et al., 2018; Deshmukh et al., 2018; Harwood et al., 2018). In sum, our results suggest that loss of PR and ER signaling in endometrial cells may lead to the aberrant expression of the genes located in TADs with PgCRs (PgCR-genes), which could contribute to tumor progression.

## Materials and Methods

### Cell culture and hormonal treatments

Endometrial adenocarcinoma Ishikawa cells and FPR Ishikawa cells were cultured in phenol red DMEM/F12 medium (GIBCO, Thermo Fisher Scientific) supplemented with 10% FCS (GreinerBioOne) and gentamycin (Thermo Fisher Scientific) at 37°C and 5% carbon dioxide to maintain cell line stock. Before each experiment, cells were plated in phenol red-free DMEM/F12 medium supplemented with 5% dextran-coated charcoal-treated (DCC)-FCS and gentamycin for 48h. Then, the medium was replaced by serum-free DMEM/F12 and kept in it for 18h (overnight). Treatments were performed with R5020 and E2 to a final concentration of 10nM and ethanol (vehicle) for the times indicated for each experiment. When indicated, pretreatment with E2 consisted of a single administration of E2 to a final concentration of 10nM 12h before hormonal treatments. T47D cells were cultured in RPMI 1640 medium as previously described (Nacht et al., 2016).

### Transfection with flag-tagged PR (FPR Ishikawa cells)

Plasmid p3xFLAG-CMV-14 carrying the complete sequence for progesterone receptor gene (HindIII924 - 938EcoRI) was introduced in Ishikawa cells using Lipofectamine 2000 (Thermo Fisher Scientific) following manufacturer recommendations. After 24h of transfection, cells were exposed to 0.6mg/ml G418 for selection. Then on, every two passages, FRP cells were exposed to a reduced concentration of G418 (0.4mg/ml), except during hormonal treatments.

### Proliferation assay

Ishikawa cells were seeded at 5×10^4^ cells/plate density in 35mm dish plates. After 48h in 5% DCC-FCS, the medium was replaced for 1% DCC-FCS for 18h. Treatments were performed for 48h and cells were then collected using trypsin (0.25%). Antagonists for ER and PR, ICI182780 and RU486 1*μ*M respectively, were added for 60min and removed before hormonal treatments. The number of live cells was determined using trypan blue (0.1%) in Neubauer chamber, repeating the procedure sixteen times for each sample and performing three independent experiments.

### BrdU incorporation assay and cell cycle analysis

Ishikawa cells were seeded and prepared for hormonal treatments as described for Proliferation assay. Treatments were carried out for 15h, the last two hours of which includes incubation with BrdU. Cells were treated with cell cycle inhibitor TSA A 250nM as negative control of BrdU incorporation. After collecting cells in trypsin and washing them with PBS, ethanol 70% was added to fix and permeabilize them. DNA denaturation was achieved with 0.5% BSA and 2M HCl after which cells were incubated in 1:2000 solution of anti-BrdU (BD Pharmingen) for 1h at RT. FITC secondary antibody (Dako) was incubated for 1h in obscurity at RT followed by propidium iodide for 5min. BrdU incorporation and cell cycle phases were evaluated by flow cytometry (BD FACS Canto II) in three replicates.

### Western blot

Cell extracts were collected at the times indicated by the experiment with 1% SDS, 25mM Tris-HCl pH 7.8, 1mM EDTA, 1mM EGTA and protease and phosphatase inhibitors. Total protein extracts were loaded in 8% SDS-PAGE and incubated with the following antibodies: PR (H190, Santa Cruz Bio.), ERalpha (HC-20, Santa Cruz Bio.) and alpha-tubulin (Sigma Aldrich). Quantification of gel images was performed with ImageJ software and expressed as abundance in relative units to alpha-tubulin.

### Immunofluorescence

Cells were seeded onto coverslips in six-well plates in a density of 10^3^ cells/ 150*μ*l using the protocol described in Cell culture and hormonal treatments and either pretreated or not with E2 10nM during the last 12h of serum-free culture. After hormonal treatments cells were washed with ice cold PBS followed by fixation and permeabilization by incubation in 70% ethanol for 12h at −20°C. After rinsing three times for 5min in 0.1% Tween-PBS, the coverslips were incubated for 2h with 10% BSA in 0.1% Tween-PBS to reduce nonspecific staining. To detect PR (H-190 Santa Cruz Bio.), phosphoserine 294 PR (S294 Cell Signaling), ERalpha (HC-20 Santa Cruz Bio.) and PAX2 (Biolegends) cells were incubated with corresponding antibodies diluted in 10% BSA 0.1%Tween-PBS at 4°C overnight. After several washes in Tween-PBS, coverslips were exposed to secondary antibodies Alexa 488 and Alexa 555 (Thermo Fisher Scientific, Thermo Fisher Scientific) diluted 1:1000 in 10% BSA 0.1% Tween-PBS for 1h at room temperature using DAPI to reveal nuclei. Coverslips were mounted on slides with Mowiol mounting medium (Sigma Aldrich) and analyzed in TIRF Olympus DSU IX83 (Olympus Life Sciences Solutions). Quantification of nuclear fluorescence was done with ImageJ software after generating a binary mask in dapi images.

### qRTPCR

After 12h of treatment with R5020 and E2, cell extracts were collected in denaturing solution (4M Guanidine thiocyanate, 25mM Sodium citrate pH 7, 0.1M 2-Mercaptoethanol, 0.5% Sarkosyl) and total RNA was prepared following phenolchloroform protocol (Chomczynski and Sacchi, 1987). Integrity-checked RNA was used to synthesize cDNA with oligodT (Biodynamics) and MMLV reverse transcriptase (Thermo Fisher Scientific). Quantification of candidate gene products was assessed by real-time PCR. Expression values were corrected by GAPDH and expressed as mRNA levels over time zero (T0). Primer sequences are available on request.

### Luciferase reporter assay

Ishikawa cells were seeded and prepared for hormonal treatments as described for Proliferation assay without addition of gentamycin. Cells were co-transfected with MMTV LTR-Firefly Luciferase (pAGMMTVLu, gift from Laboratory of Patricia Elizalde) and CMV-Renilla luciferase (pRL-CMV, Promega) plasmids using lipofectamine plus 2000 (Thermo Fisher Scientific). After 5h, media were renewed with the addition of antibiotics and 12h later cells were treated with vehicle (ethanol) and R5020 for 20h. Firefly and Renilla activities (arbitrary units) were determined with Dual-Luciferase Reporter assay system (Promega) and expressed as Firefly units relative to internal control Renilla for each sample (Firefly x10^4^/Renilla).

### RNAseq

Total RNA was collected from untreated (T0) and 12h R5020- and E2-treated Ishikawa cells using RNeasy Plus Mini Kit (QIAGEN) and subjected to high-throughput sequencing in Illumina HiSeq 2000 and 2500. Poly-A-enriched RNA was used to prepare libraries with TruSeq RNA Sample Preparation kit v2 y v4 (ref. RS-122-2001/2, Illumina) according to instructions from manufacturer followed by single-end (run1) and paired-end (run2) sequencing. Good quality 50bp reads were aligned to the reference human genome (hg19, UCSC) using Tophat software (Trapnell et al., 2009) keeping those that mapped uniquely to the reference with up to two mismatches and transcript assembly, abundance quantification and differential expression analyses were performed with the Cufflinks tool (Trapnell et al., 2010). Genes under 200bp in length or with FPKM values below 0,1 were excluded from downstream analyses. Genes were classified into induced, repressed or non-regulated depending on log2FC value relative to untreated cells (T0). Threshold value was arbitrarily set at log2FC = ± 0.8 and q<0.05 (FDR). Enriched terms and TFBS were determined through RDAVIDWebservice (Fresno and Fernandez, 2013) and DAVID web-based tool (Huang et al., 2009) under standard parameter settings for each tool.

### Gene Set Enrichment Analysis (GSEA)

GSEA tool was implemented following instructions from developers under default parameters (Subramanian et al., 2005). The expression dataset was created using Ishikawa RNAseq results, labelling samples as “R5020” and “E2” for categorical classification (phenotypes). Gene sets were constructed from proliferative (SRR9298724, SRR9298725, SRR9298726 and SRR9298727) and mid-secretory (SRR9298728, SRR9298729, SRR9298730, SRR9298731 and SRR9298732) normal endometrial RNAseq samples (Chi et al., 2020). Differential expression analysis to extract genes representative of each stage was performed with DESeq2 package (|log2FC|>2.5, p<0.05) (Love et al., 2014).

### Endometrial cancer samples (TCGA)

Raw count data from endometrial cancer RNAseq samples (n=575) were downloaded from The Cancer Genome Atlas (TCGA), project TCGA-UCEC. Endometrioid adenocarcinoma samples (n=423) were selected using associated clinical data and only protein coding genes above arbitrary threshold (mean > 100 counts) were kept for further analyses. Raw counts were normalized in DESeq2 package and later used for heatmaps (pheatmap R package (Kolde and Kolde, 2015)) and Principal Component Analysis in PCAtools package (Blighe et al., 2019).

### Chromatin immunoprecipitation (ChIP)

ChIP experiments were performed as described in (Strutt and Paro, 1999) and (Vicent et al., 2011). Antibodies used for immunoprecipitation were PR (H190, Santa Cruz Bio.), ERalpha (HC-20X and H184X, Santa Cruz Bio.), PAX2 (PRB-276P, BioLegend) and normal rabbit IgG (sc-2027, Santa Cruz Bio.). Enrichment to DNA was expressed as percentage of input (non-immunoprecipitated chromatin) relative to untreated Ishikawa cells (T0) using the comparative Ct method. Ct values were acquired with BioRad CFX Manager software.

### ChIPseq

After minor modifications to the ChIP protocol described in (Vicent et al., 2011), purified ChIP-DNA was submitted to deep sequencing using Illumina HiSeq-2000. Libraries were prepared by the Genomics unit of the CRG Core Facility (Centre for Genomic Regulation, Barcelona, Spain) with NEBNext ChIPseq Library Prep Reagent Set (ref. E6200S, Illumina) and 50bp sequencing reads were trimmed to remove Illumina adapters and low-quality ends using Trimmomatic (Bolger et al., 2014) version 0.33 in single-end mode. Good quality reads were aligned to the reference human genome (hg19, UCSC) with BWA (Li and Durbin, 2009) v0.7.12 (BWA_MEM algorithm with default parameters) keeping alignments that mapped uniquely to the genome sequence (Samtools version 1.2, (Li and Durbin, 2009)). Overlapping reads were clustered and significant signal enrichments (peaks) were identified by MACS2 v2.1.0 (Zhang et al., 2008) using input as background signal. FDR value during initial peak calling steps was set to 0.05 (q), though downstream analyses included only those with q<10^-5^. Replication of binding sites was evaluated among treatments (time of exposure to hormone) and conditions (no pretreated, pretreated and FPR) using scatter plots and venn diagrams. Selected sites were validated by qPCR. When necessary peak files were converted to hg38 coordinates using the batch conversion tool from UCSC. ChIPseq coverage data of proliferative and secretory normal endometrium were downloaded from GEO (GSE132713, (Chi et al., 2020)).

### Heatmaps, Scatterplots and Motif analysis

Overlap of ChIPseq peak regions defined by upstream peak calling procedures (MACS2) were determined using intersectBed program from the bedTools suite (Quinlan, 2014). An overlap of at least one bp was considered positive. De novo motif discovery (MEME software) performed on sequences contained in 10kb windows centered in peak summits. Graphs, correlation tests, non-linear regression and statistical analyses in general were performed for common peaks between ChIPseq samples using R (R Development Core Team). Heatmaps were plotted using the summit of the peaks as a reference central position. Reference positions were taken from common and exclusive peaks within experiments and were sorted by height of the peak. Genome aligned reads occurring between −5000 and +5000 bp from reference sites were mapped using count_occurences program (Kremsky et al., 2015) and the number of reads per bins of 200bp was used for the color intensity of heatmap cells with R. For Motif discovery, genomic regions of top 500 peaks ranked by their height were extracted from each set and regions that overlap with repeats, low complexity regions or transposable elements (extracted from the UCSC genome browser, hg19 human assembly), were removed from the analysis. Motif discovery was performed using MEME program suite executed with the following parameters: −maxsize 250000 - revcomp −dna −nmotifs 3 −mod oops (Bailey et al., 2015). Motif enrichments were evaluated with the procedure and statistics described in (Agirre et al., 2015). Additionally, the analysis utilized a 5mers collection of 1,395 human position frequency matrices modelling transcription factors binding sites (Weirauch et al., 2014), which were scanned (p-value<1e^-4^) and their enrichment evaluated in regions of 200bp centered in the summits of whole peaks sets. To uncover motif profiles, discovered and library motifs were wholegenome scanned (p-value<1e^-4^). Their occurrences around the sets of summits were obtained with count-occurrences (±2000bp, bin size=200bp) and the profiles showing the proportion of regions per bin having at least one match were plotted using R.

### Binding site-gene association

Genomic coordinates of PR and ERalpha binding sites (hg38) were fed to GREAT web tool (McLean et al., 2010) to identify potential cis-regulatory interactions. Association was determined in a “basal plus extension” process using a proximal regulatory domain of 5kb upstream and 1kb downstream from each TSS (GRCh38, UCSC hg38) and an extension of 100kb in both directions. The group of genes associated with PRbs or ERbs were respectively intersected to R5020 and E2 RNAseq results, employing simple python scripting.

### ATACseq

ATACseq was performed as previously described (Buenrostro et al., 2013). Briefly, 50,000 cells were lyzed with 50*μ*l cold lysis buffer (Tris-Cl pH 7.4 10mM; NaCl 10mM; MgCl2 3mM; NP-40 0.1% v/v) and centrifuged at 500xg for 10min at 4°C. Nuclei were resuspended in TD Buffer with 1.5*μ*l Tn5 Transposase (Nextera, Illumina) and incubated 15 minutes at 37^°^C. DNA was isolated using Qiagen MinElute column and submitted to 10 cycles of PCR amplification using NEBNext High-Fidelity 2X PCR Master Mix (Univ. primer: AATGATACG-GCGACCACCGAGATCTACACTCGTCG-GCAGCGTCAGATGTG; Indexed primers: CAAGCAGAAGACGGCATACGA-GATNNNNNNNNGTCTCGTGGGCTCG-GAGATGT). Library were size selected using AMPure XP beads and sequenced on a NextSeq 500 instrument (2×75nt).

### Hi-C

High-throughput chromosome conformation capture assays were performed as previously described (Lieberman-Aiden et al., 2009; Rao et al., 2014). Adherent cells were directly cross-linked on the plates with 1% formaldehyde for 10min at room temperature. After addition of glycine (125mM final) to stop the reaction, cells were washed with PBS and recovered by scrapping. Cross-linked cells were incubated 30min on ice in 3C lysis Buffer (10mM Tris-HCl pH=8, 10mM NaCl, 0.2% NP40, 1X anti-protease cocktail), centrifuged 5min at 3,000 rpm and resuspended in 190*μ*l of NEBuffer2 1X (New England Biolabs - NEB). 10*μ*l of 10% SDS were added and cells were incubated for 10min at 65°C. After addition of Triton X-100 and 15min incubation at 37^°^C, nuclei were centrifuged 5min at 3,000 rpm and resuspended in 300*μ*l of NEBuffer2 1X. Digestion was performed overnight using 400U MboI restriction enzyme (NEB). To fillin the generated ends with biotinylated-dATP, nuclei were pelleted and resuspended in fresh repair buffer 1x (1.5*μ*l of 10mM dCTP; 1.5*μ*l of 10mM dGTP; 1.5*μ*l of 10mM dTTP; 37.5*μ*l of 0.4mM Biotin-dATP; 50U of DNA Polymerase I Large (Klenow) fragment in 300*μ*l NEBuffer2 1X). After 45min incubation at 37^°^C, nuclei were centrifuged 5min at 3,000 rpm and ligation was performed 4h at 16°C using 10,000 cohesive end units of T4 DNA ligase (NEB) in 1.2ml of ligation buffer (120*μ*l of 10X T4 DNA Ligase Buffer; 100*μ*l of 10% Triton X-100; 12*μ*l of 10mg/mL BSA; 963*μ*l of H2O). After reversion of the cross-link, DNA was purified by phenol extraction and EtOH precipitation. Purified DNA was sonicated to obtain fragments of an average size of 300-400bp using a Bioruptor Pico (Diagenode; 8 cycles; 20s on and 60s off). 3*μ*g of sonicated DNA was used for library preparation. Briefly, biotinylated DNA was pulled down using 20*μ*L of Dynabeads Myone T1 streptavidine beads in Binding Buffer (5mM Tris-HCl pH7.5; 0.5mM EDTA; 1M NaCl). End-repair and A-tailing were performed on beads using NEBnext library preparation endrepair and A-tailing modules (NEB). Illumina adaptors were ligated and libraries were amplified by 8 cycles of PCR. Resulting Hi-C libraries were first controlled for quality by low sequencing depth on a NextSeq500 prior to higher sequencing depth on HiSeq2000. Hi-C data were processed using an in-house pipeline based on TADbit (Serra et al., 2017). Reads were mapped according to a fragment-based strategy: each side of the sequenced read was mapped in full length to the reference genome Human Dec. 2013 (GRCh38/hg38). In the case reads were not mapped when intra-read ligation sites were found, they were split. Individual split read fragments were then mapped independently. We used the TADbit filtering module to remove non-informative contacts and to create contact matrices as previously described (Serra et al., 2017) PCR duplicates were removed and the Hi-C filters applied corresponded to potential non-digested fragments (extra-dandling ends), non-ligated fragments (dandling-ends), self-circles and random breaks.

### CNV

The copy number variation (CNV) analysis was estimated comparing the coverage obtained in the Hi-C datasets with the expected coverage for a diploid genome based on the density of restriction sites and genomic biases (Vidal et al., 2018). Indeed, the linear correlation between number of Hi-C contacts and number of restriction sites is lost in case of altered copy number allowing the estimation of a relative number of copy as compared to diploid chromosomes in each dataset. Such estimations are consistent with other analyses and with karyotyping (Le Dily et al., 2014).

### Virtual 4C

Hi-C matrices were normalized for sequencing depth and genomic biases using OneD (Vidal et al., 2018) and further smoothed using a focal average. Virtual 4C plots were generated from the matrices locally normalized and expressed as normalized counts per thousands within the region.

### Intra-TAD interactions between specific loci

Each bin of a TAD was labeled as part of a PgCR or TSS (or “others” if they did not belong to the previous types). We collected the observed contacts between the different types of bins and computed the expected contacts frequencies based on the genomic distance that separate each pair. In the figure, results are expressed as Log2 of the ratio of observed contacts between the different types of pairs above the intra-TAD background.

## Supporting information

PgCR_coordinates

## Data Availability

All raw and processed sequencing data generated in this study have been submitted to the NCBI Gene Expression Omnibus (GEO; https://www.ncbi.nlm.nih.gov/geo/) under accession number GSE139398.

## Acknowledgments

We are grateful to members of the Beato and Saragüeta laboratories for help and suggestions.

## Author Contributions

ALG, FLD, MB and PS design experiments. ALG, FLD, RJ, GV and ITR performed cell culture and experiments. ALG, NB, RJ, GM, CF, JQO, EV and FLD performed bioinformatic analyses. RJ, JLV, EV and FLD analyzed Hi-C results. GV, EF, GPV, MB, ALG and PS discussed experiments and manuscript. MB and PS provided fundings for this paper. ALG, MB and PS wrote the manuscript.

