## Supplemental Figures for "Higher-order chromatin organization defines Progesterone Receptor and PAX2 binding to regulate estradiol-primed endometrial cancer gene expression"


#### Running title

Endometrial hormone-dependent PR gene regulation.

### Supplemental Material

La_Greca_Supplemental_FigS1

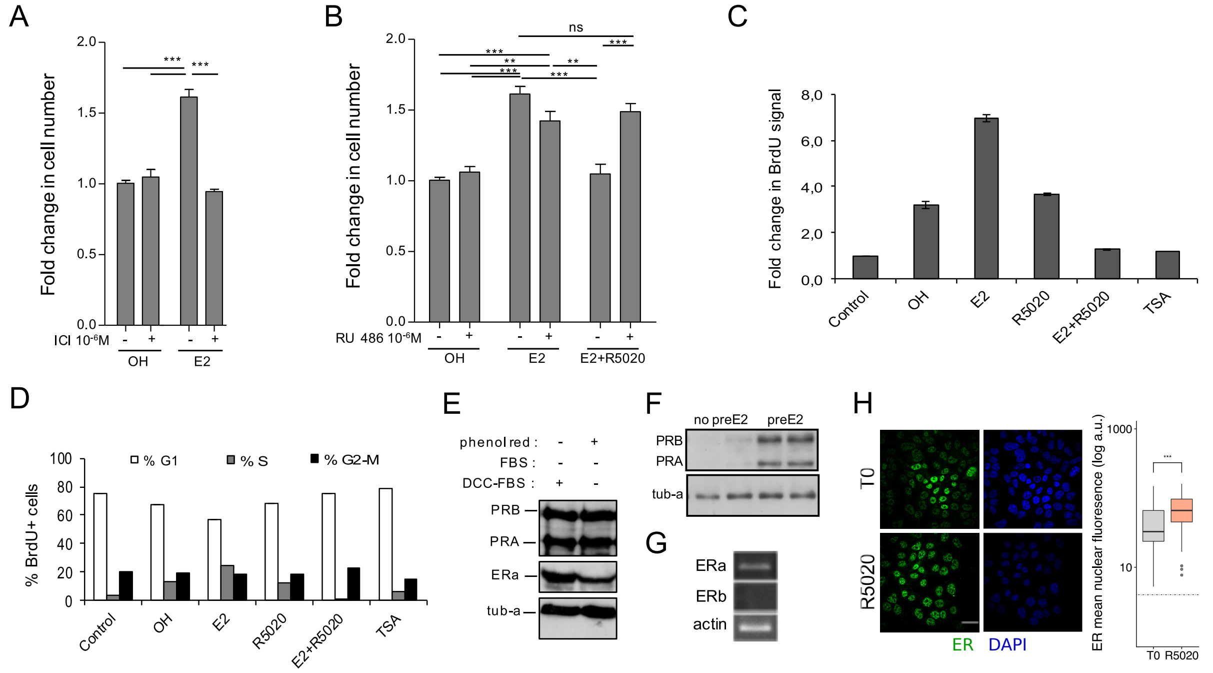


**Supplementary Figure S1. E2-dependent cell cycle progression and proliferation of Ishikawa cells is mediated by ERalpha. R5020 antagonism on E2 is mediated by PR.** (A) Cells were pretreated with ICI182780 1$\mu$M (ICI $10^{-6}$M) and then treated with vehicle (OH) and E2 10nM (E2) for 48h. Number of cells was determined and presented as in Figure 1A. (***) p<0,001. (B) Cells were pretreated with RU486 1$\mu$M (RU 486 $10^{-6}$M) and later treated with vehicle (OH), E2 10nM (E2) and E2 and R5020 (E2+R5020) for 48h. Number of cells was determined and expressed as in A. (**) p<0,01; (***) p<0,001; ns: not significant. (C) BdrU incorporation was measured in untreated cells (control) and treated with vehicle (OH), E2 10nM (E2), R5020 10nM (R5020), E2 and R5020 (E2+R5020) and TSA 250nM (TSA) for 15-18h. Mean fold change in BrdU positive cells for each treatment over Control ± SE of three independent experiments is shown. (D) Percentage of BdrU positive cells in each phase of the cell cycle for treatments shown in D: %G1 (white bars), %S (grey bars) and %G2-M (black bars). (E) PR (PRB and PRA) and ERalpha (ERa) protein levels of serum-deprived Ishikawa cells were assayed by western blot (Ishikawa) in two different conditions: phenol red-positive medium (phenol red +) and phenol red-negative medium (phenol red -) with 5% dextran-charcoal-treated serum (DCC-FBS). Alfa tubulin was used as loading control (tub-a). Western blot image at the bottom of the figure shows PR protein levels in non-pretreated (no preE2) and 12h E2-pretreated (preE2) Ishikawa cells using alfa tubulin as loading control (tub-a). (F) Western blot image showing PR protein levels in Ishikawa cells pretreated with E2 for 12h (preE2) and non-pretreated (no preE2). Alfa tubulin was used as loading control (tub-a). (G) mRNA levels of ERalpha, ER$\beta$ and actin in serum-deprived Ishikawa cells. (***) p<0.001. (H) Representative image of anti-ERalpha detection in Ishikawa cells treated or not with R5020 for 60min. Quantification was performed as in Figure 1. Nuclei were revealed with DAPI (blue).

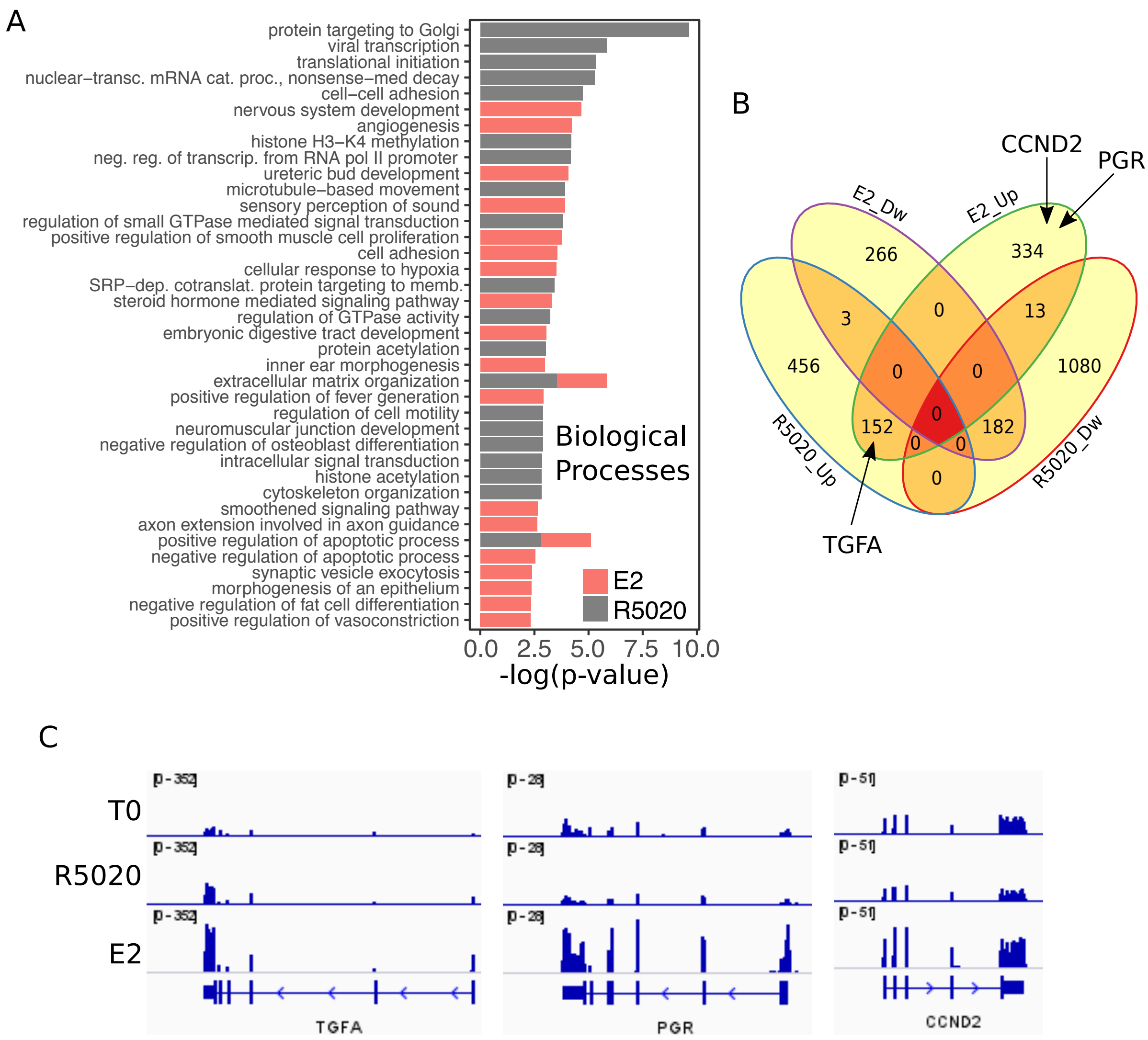


**Supplementary Figure S2. RNAseq results from hormone-treated Ishikawa cells.** (A) Overrepresented Biological Processes in RNAseq results from Ishikawa cells treated with R5020 and E2 for 12h analysed using DAVID web-based tool. Only the top 20 terms with $p<0.05$ were selected for each hormone. (B) Venn Diagram represents intersection between differentially expressed genes (v.T0; $\log2FC=\pm0.8$, $q<0.05$) in Ishikawa cells treated with R5020 and E2 for 12h. Upregulated and downregulated genes are indicated by Up and Dw, respectively. (C) Normalized expression profiles of genes located in the proximity of steroid receptor binding. RNAseq results expressed in RPKM for genes *TGFA*, *PGR* and *CCND2* displayed in IGV-based environment. Tracks for untreated (T0) and R5020 10nM (R5020) and E2 10nM (E2) 12h treatments were represented in the same vertical viewing range value.

La_Greca_Supplemental_FigS3

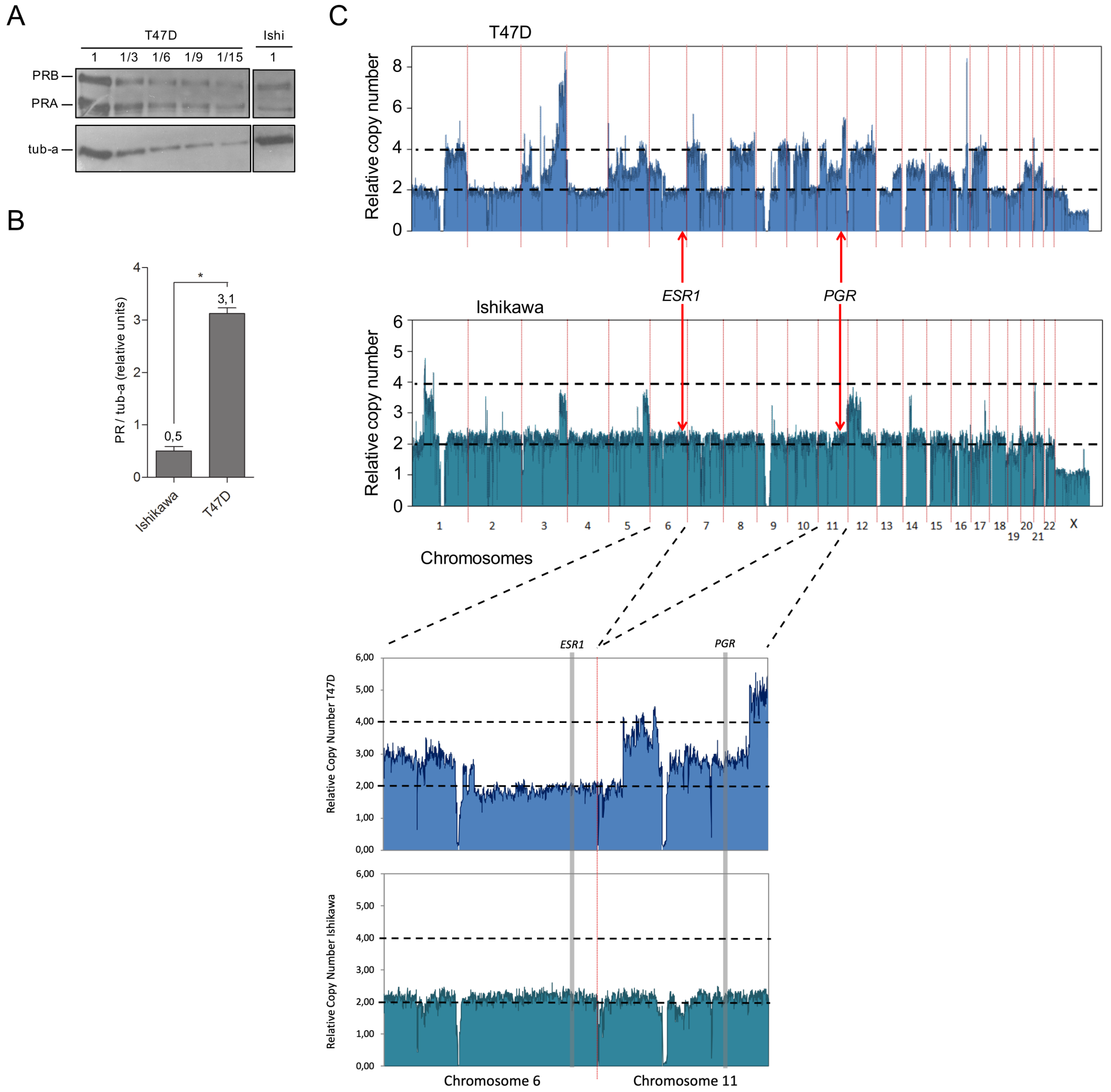


**Supplementary Figure S3. Ishikawa cells express six times less PR than T47D cells.** (A) Representative image of western blot assays using protein extracts from Ishikawa and T47D cells. 30$\mu$g (1) of T47D extracts were seeded followed by 10$\mu$g (1/3), 5$\mu$g (1/6), 3.3$\mu$g (1/9) and 2$\mu$g (1/15) of the same extract. 30$\mu$g of protein extract from Ishikawa cells (Ishi) were loaded on the same gel. PR isoforms (PRA and PRB) levels were evaluated and tubulin (tub-a) as control. Image was processed to remove lanes that did not contribute to the present analysis. (B) Quantification of PR protein levels in Ishikawa and T47D cells expressed as mean units of PRA+PRB relative to tubulin protein levels (tub-a) of three independent experiments. Numbers above bars indicate value of quantification relative to tubulin determined by densitometry of western blot assays. (*) $p<0{,}05$. (C) Copy number variation (CNV) analysis of 100kb regions distributed over all chromosomes (hg19 UCSC) in T47D (upper panel) and Ishikawa (lower panel) cell lines. Black dotted lines limit diploid (2) and tetraploid (4) numbers. Location of estrogen receptor 1 (ESR1) and progesterone receptor (PGR) loci are indicated with red arrows. Enlarged image of regions covering chromosome 6 (*ESR1*) and chromosome 11 (*PGR*) is shown below graph.

La_Greca_Supplemental_FigS4

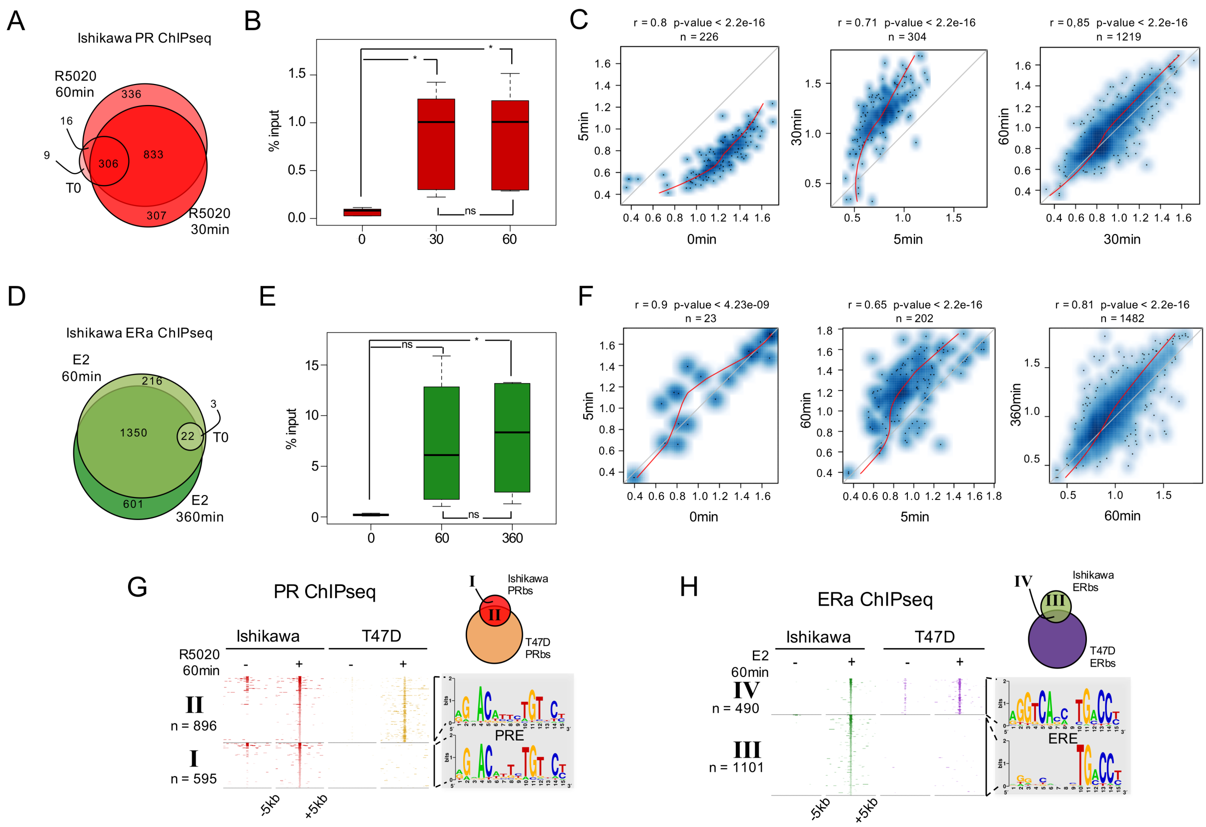


**Supplementary Figure S4. Genome-wide analysis of PR and ERalpha binding in Ishikawa cells.** (A) ChIPseq results using anti-PR antibody in Ishikawa cells. Cells were treated or not (T0) for 5min (R5), 30min (R30) and 60min (R60) with R5020 10nM. The number of PRbs for T0, R30 and R60 is indicated in the Venn Diagram. (B) Boxplot shows validated PR binding sites expressed as percentage of input (%input) for 0, 30 and 60min R5020-treated Ishikawa cells. Sites included in this analysis are located in: promoter of ALPP gene, intergenic sequence upstream of ALPP gene, intergenic sequence upstream of EGFR gene, distal promoter of SERPINA3 gene, promoter of RPS6KA1 and intragenic sequence in TGFA gene. Significance was evaluated by ANOVA followed by Tukey's multiple contrasts and significant results are indicated for each individual comparison. (ns) not significant; (*) p<0,05. (C) Scatterplots depict correlation in peak signal intensity among samples. Number of sites in common (n) as wells as correlation coefficient (r) and p-value are shown over each plot. Non-linear regression curves are indicated as red lines. (D) ChIPseq results using anti-ERalpha antibody in Ishikawa cells treated or not (T0) with E2 10nM for 5 (E5), 60 (E60) and 360min (E360). In the Venn Diagram are shown untreated (T0), E60 and E360 as well as the number of ER binding sites (ERbs) for each group. (E) Boxplots shows validated ERalpha binding sites expressed as percentage of input (%input) for 0, 60 and 360min E2-treated Ishikawa cells. Sites included in this analysis are located in: intergenic region upstream of ALPP gene, intergenic region upstream of TGFA gene, intragenic region upstream of TGFA gene and promoter of TIPARP gene. Significance was evaluated by ANOVA followed by Tukey multiple contrast tests and significant results are indicated for each individual comparison. (ns) not significant; (*) p<0,05. (F) Scatterplots as in C depict correlation in peak signal intensity among samples immunoprecipitated with anti-ERalpha. (G) Heatmaps of PR ChIPseq data from untreated (-) and 60min R5020-treated (+) Ishikawa and T47D cells. Regions were defined inside a window of -5kb to +5kb centered in peak summit. Intensity of the signal corresponds to the number of reads in the region. ChIPseq data are organized according to the Venn diagram scheme on the top right side of the figure as follows: I, PRbs solely found in Ishikawa cells; II: PRbs shared by both cell lines. De novo discovered motifs in I and II are indicated as sequence logos to the right of the map. PRE: progesterone response element. (H) Heat map constructed as in G of ERalpha ChIPseq data from untreated (-) and 60min E2-treated (+) Ishikawa and T47D cells. ChIPseq data are organized according to the Venn diagram scheme on the top right site of the figure as follows: III, ERbs solely found in Ishikawa cells; IV: ERbs shared by both cell lines. Discovered motifs in III and IV indicated presented as sequence logos are on the right site of the map. ERE: estrogen response element.

La_Greca_Supplemental_FigS4

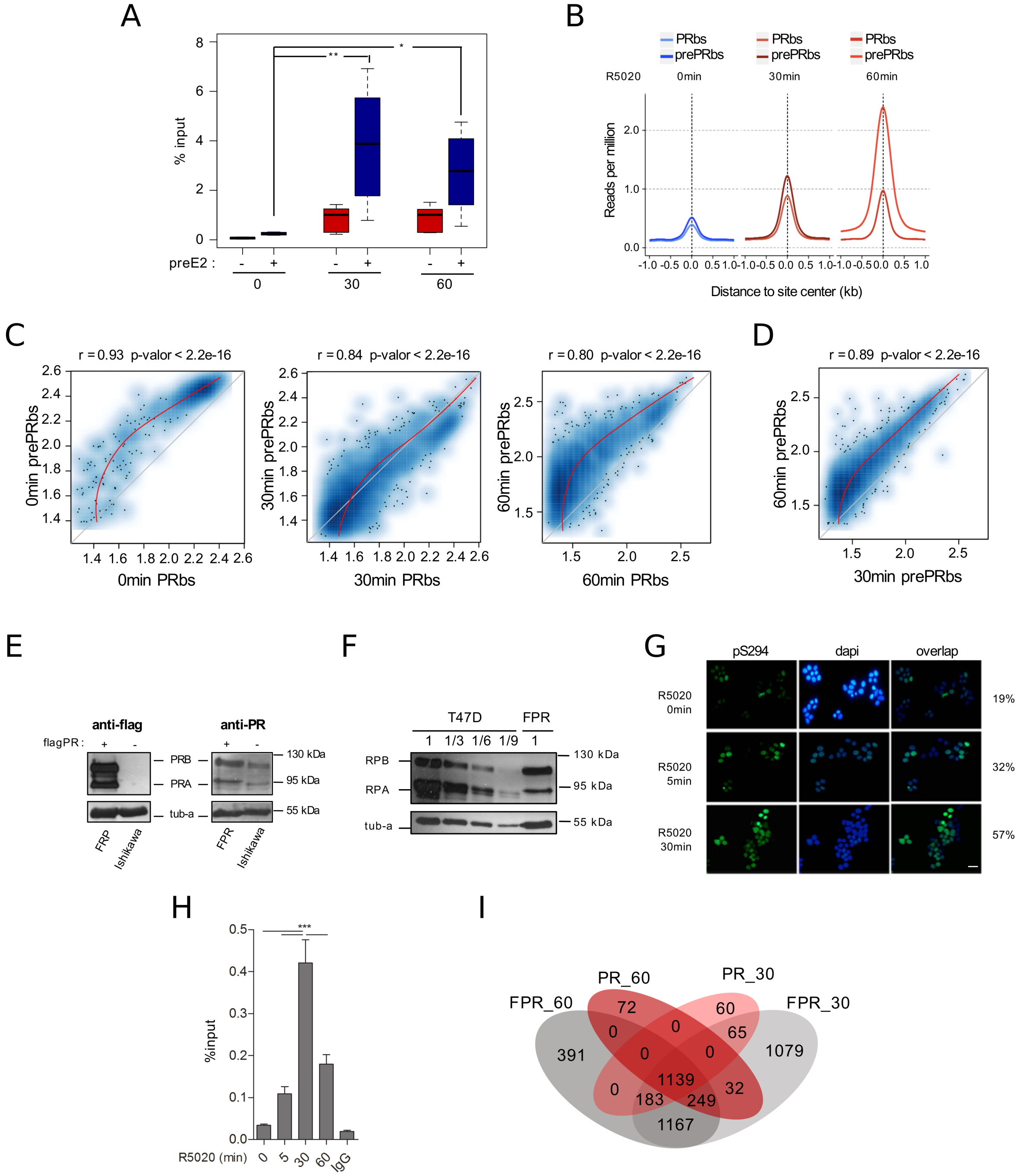


**Supplementary Figure S5. PR overexpression on Ishikawa cells.** (A) Validation of PR binding sites in non-pretreated (preE2 -) and 12h E2-pretreated (preE2 +) Ishikawa cells expressed as percentage of input (%input) for 0, 30 and 60min treatments with R5020. Regions included in the analysis are described in Supplementary Figure S4B. Significance was evaluated by ANOVA followed by Tukey multiple contrast tests and significant results are indicated for each individual comparison. (ns) not significant; (*) p<0,05; (**) p<0,01. (B) Signal strength for PR and preE2PR binding of untreated (T0) and 30 and 60min R5020-treated cells. Intensity was defined as reads per million relative to distance to site center in kb inside a window of -/+ 1kb. (C) Correlation in PR peak signal intensity between 12h E2-pretreated (prePRbs) and non-pretreated (PRbs) Ishikawa cells treated or not (0min) with R5020 (30 and 60min). Correlation coefficient (r) and p-value are shown over each plot. Non-linear regression curves are indicated as red lines. (D) Correlation between 30min and 60min R5020-treated prePRbs. (E) Western blot corroboration of PR protein levels on Ishikawa and FPR Ishikawa (exogenous expression of triple flag-PR) cells. Antibodies used in the analysis detected the flag tag (anti-flag) or PR (PRA and PRB, anti-PR). Tubulin (tub-a) was used as normalization control. (F) Representative image of western blot assays using protein extracts from FPR Ishikawa and T47D cells. 30$\mu$g (1) of T47D extract were seeded followed by 10$\mu$g (1/3), 5$\mu$g (1/6), 3.3$\mu$g (1/9) and 2$\mu$g (1/15) of the same extract. 30$\mu$g of protein extract from FPR Ishikawa cells (FPR) were loaded on the same gel. Anti-PR antibody was used to reveal PR isoforms (PRA and PRB) and tubulin (tub-a) as control. Lanes that did not contribute to the present analysis were removed. (G) Immunofluorescence evaluation of serine 294 phosphorylated PR (pS294) is shown in the first column (green dots). Nuclei were dyed with dapi (blue dots) (second column). Overlap of green and blue signals is shown in the third column. Percentage of positive nuclear pS294 Ishikawa cells (first column) was calculated as green dots x 100/blue dots in untreated cells (T0) and after treating them for 5 (T5) and 30min (T30) with R5020 10nM. (H) Recruitment of PR to the EGFR enhancer sequence -described in Figure 1F- in FPR cells treated for 0, 5, 30 and 60min with R5020 10nM and unspecific IgG control. Results are expressed as in Figure 1F (two independent experiments). (***) p<0,001. (I) Venn Diagram shows shared binding sites among 30min- and 60min-treated FPR (FPR_30 and FPR_60) and parental Ishikawa cells (PR_30 and PR_60).

La_Greca_Supplemental_FigS6

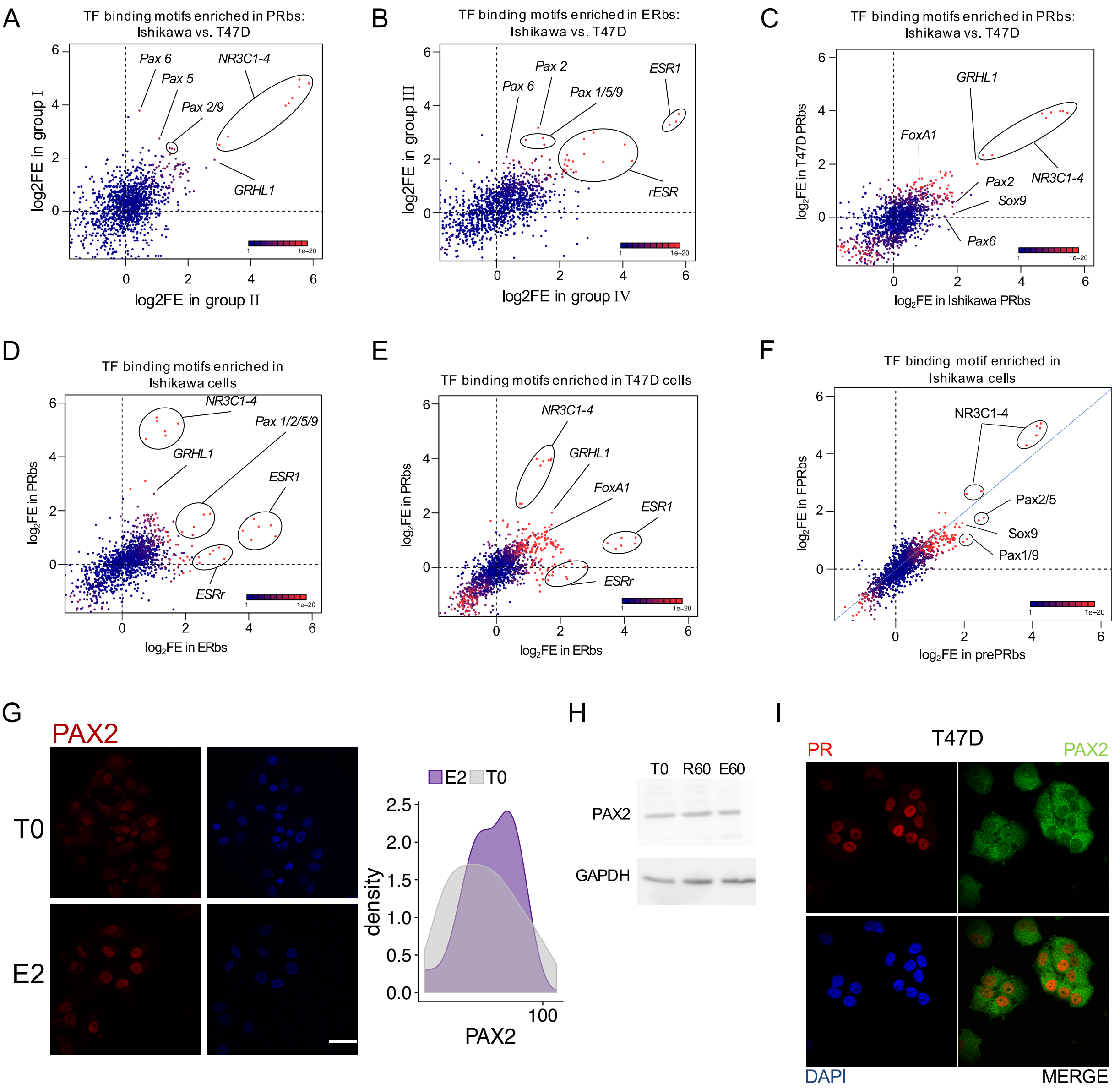


**Supplementary Figure S6. PAX2 binding to chromatin is hormone dependent in Ishikawa cells while it is not localized to nuclei of T47D cells.** Scatter plots showing fold enrichment values (log2FE) of 1,395 known TF binding motifs on (A) PRbs in group I and group II from Supp Fig S4G, (B) ERbs in group III and group IV from Figure S4H, (C) PRbs from Ishikawa and T47D cells, (D) ERbs and PRbs from Ishikawa cells, (E) ERbs and PRbs from T47D cells and (F) FPRbs and prePRbs. Combined p-values (product of x and y p-value multiplication) for enrichment analyses are indicated through the color key displayed at the lower right corner of the plots. Relevant motifs pointed on the plots correspond to NR3C1-4, ESR1, rESR, members of the Pax family (1, 2, 5, 6 and 9), GRHL1 (TF involved in epithelial development), SOX9 and FOXA1. (G) Representative images of PAX2 signal (red) in cells treated or not with E2 for 60min and dapi staining (blue). Scale bar is shown in the figure and equals to 30$\mu$m. Distribution of intensities in nuclear signal for both conditions is shown to the right of the images. (H) Western blot image of PAX2 protein levels in untreated (T0) Ishikawa cells and treated with R5020 for 60min (R60) or E2 for 60min (E60). GAPDH was used as normalizing control. (I) PR (red) and PAX2 (green) staining in T47D cells treated with R5020 for 60min. Dapi was used for revealing nuclei.

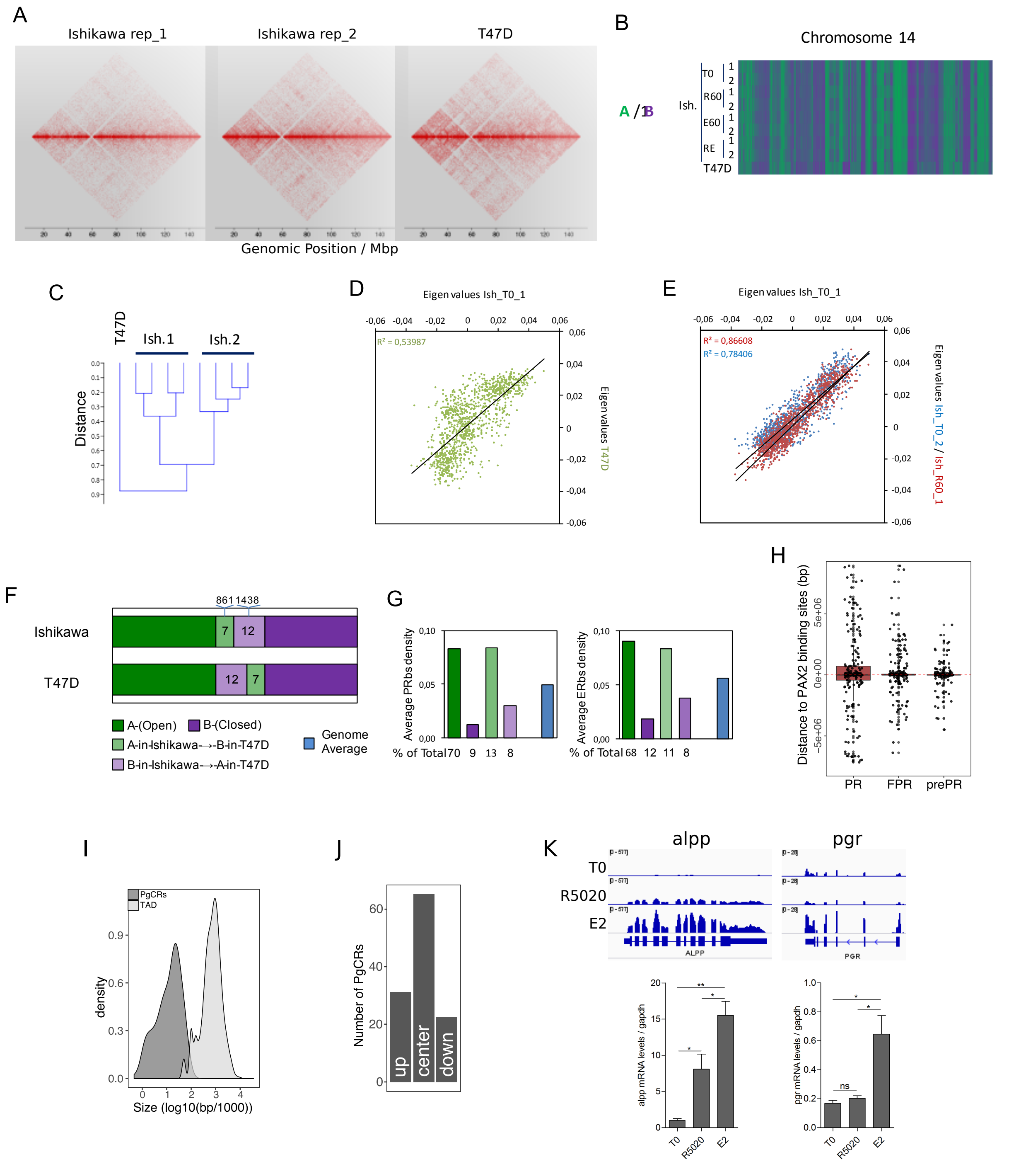


**Supplementary Figure S7. Chromosome compartments analysis and definition of Progesterone Control Regions.** (A) Contact matrices of chromosome 12 region at 100kb resolution obtained by In Nucleo Hi-C performed in two replicates of Ishikawa (Ishikawa_rep1 and Ishikawa_rep2) and T47D cells. (B) Hi-C datasets of untreated (T0) Ishikawa cells or treated with R5020 (R60), E2 (E60) and both hormones combined (RE60) were used to determine the spatial segregation of chromatin in open (A: green) or closed (B: purple) chromatin compartments compared to T47D cells. A fraction of chromosome 14 is shown. (C) Relationship among samples was evaluated after clustering Hi-C datasets. Plot reflects differences as a function of distance. (D and E) Correlation analysis between Ishikawa replicates and T47D samples using eigen values determined from Hi-C datasets. Correlation coefficients are indicated in the figures. (F) Genome-wide proportions of 100kb windows of the genome found commonly in A (green) or B (purple) in the two cell lines or showing divergent chromatin states (light green and light purple). (G) Histograms showing the average density of PR and ER binding sites (per 100kb) in the different types of regions mentioned in F. Below are the proportion of total PR and ERalpha binding sites found in those distinct regions. (H) Relative distance in bp from PRbs, FPRbs and prePRbs to PAXbs (both E2 and R5020 originated sites). Red dashed line marks overlapping events (distance = 0). Peaks located upstream of PAXbs are noted as negative distance values, while downstream peaks are positive. (I) Size distribution of PgCRs and TADs, expressed as log10. (J) Distribution of PgCRs in harbouring TADs divided into center (center), upstream end (up) and downstream end (down). (K) RNAseq results for genes *ALPP* and *PGR* displayed in a genome browser-based environment. Tracks for untreated (T0) and R5020 10nM (R5020) and E2 10nM (E2) 12h treatments were represented in the same vertical viewing range value (vvrv). A different vvrv was used for each gene in display. Quantitative PCR validations of RNAseq results are shown below. Results are expressed as mRNA levels/gapdh ± SE of three independent experiments. (ns) not significant; (*) p<0,05; (**) p<0,01.
