## Supplementary material for "Higher-order chromatin organization defines Progesterone Receptor and PAX2 binding to regulate estradiol-primed endometrial cancer gene expression": PgCR_coordinates

| | | |
|---|---|---|
| chr1 | 14976339 | 14977774 |
| chr1 | 61424138 | 61447183 |
| chr1 | 78003875 | 78005040 |
| chr1 | 107738782 | 107748568 |
| chr1 | 162223966 | 162235856 |
| chr1 | 172351497 | 172357060 |
| chr1 | 198598545 | 198623120 |
| chr1 | 198755914 | 198764119 |
| chr10 | 59793978 | 59808081 |
| chr10 | 59962355 | 59996509 |
| chr11 | 15657163 | 15664453 |
| chr11 | 30422270 | 30424338 |
| chr11 | 101261964 | 101299985 |
| chr11 | 130695669 | 130728760 |
| chr12 | 31721926 | 31786075 |
| chr12 | 42643653 | 42670048 |
| chr12 | 65308769 | 65309707 |
| chr12 | 69085827 | 69108471 |
| chr12 | 74941728 | 74946218 |
| chr12 | 97556021 | 97576288 |
| chr13 | 33369283 | 33404976 |
| chr13 | 69921163 | 69955568 |
| chr13 | 100398267 | 100400499 |
| chr14 | 25078122 | 25092731 |
| chr14 | 32679708 | 32714341 |
| chr14 | 53850771 | 53873410 |
| chr14 | 68168268 | 68173129 |
| chr15 | 33110847 | 33135846 |
| chr15 | 65295842 | 65305448 |
| chr15 | 71495159 | 71550256 |
| chr15 | 74642193 | 74664723 |
| chr15 | 92705707 | 92725668 |
| chr17 | 43434368 | 43457597 |
| chr17 | 71631422 | 71636192 |
| chr18 | 3538572 | 3542181 |
| chr18 | 22821854 | 22872276 |
| chr18 | 26440590 | 26511995 |
| chr18 | 33168928 | 33184521 |
| chr2 | 67144944 | 67166140 |
| chr2 | 67894983 | 67899738 |
| chr2 | 70596678 | 70616658 |
| chr2 | 102114949 | 102132197 |
| chr2 | 146615780 | 146625380 |
| chr2 | 159062908 | 159091867 |
| chr2 | 182816770 | 182840542 |
| chr2 | 182890178 | 182916916 |
| chr2 | 209472595 | 209496779 |
| chr2 | 211396713 | 211432631 |
| chr2 | 232374896 | 232378773 |
| chr2 | 234241852 | 234244245 |
| chr20 | 10861250 | 10867647 |
| chr3 | 29466132 | 29486488 |
| chr3 | 30334318 | 30371205 |
| chr3 | 79552491 | 79592056 |
| chr3 | 79651372 | 79653712 |
| chr3 | 79917601 | 79940610 |
| chr3 | 89787606 | 89796030 |
| chr3 | 169244154 | 169306122 |

| | | |
|---|---|---|
| chr4 | 4481197 | 4522369 |
| chr4 | 18930444 | 18939130 |
| chr4 | 19738900 | 19776495 |
| chr4 | 30757137 | 30789884 |
| chr4 | 73675498 | 73705272 |
| chr4 | 76517237 | 76563933 |
| chr4 | 85092612 | 85095373 |
| chr4 | 87117394 | 87130976 |
| chr4 | 138112756 | 138199982 |
| chr4 | 156763231 | 156779520 |
| chr4 | 156906167 | 156949890 |
| chr5 | 20353337 | 20360765 |
| chr5 | 31106093 | 31121800 |
| chr5 | 36422787 | 36451095 |
| chr5 | 56356266 | 56357814 |
| chr5 | 57803482 | 57825740 |
| chr5 | 92145891 | 92148092 |
| chr5 | 103724211 | 103744491 |
| chr5 | 103867749 | 103924798 |
| chr5 | 123809362 | 123814803 |
| chr5 | 124649442 | 124662954 |
| chr5 | 124736519 | 124746837 |
| chr5 | 125175268 | 125176596 |
| chr6 | 6678627 | 6726217 |
| chr6 | 11364974 | 11412953 |
| chr6 | 11472807 | 11479659 |
| chr6 | 18457103 | 18504095 |
| chr6 | 35590600 | 35602434 |
| chr6 | 74741667 | 74774866 |
| chr6 | 75333361 | 75351828 |
| chr6 | 122033436 | 122099011 |
| chr6 | 130902877 | 130908592 |
| chr6 | 136058831 | 136068248 |
| chr6 | 139973979 | 139979815 |
| chr7 | 12706624 | 12716343 |
| chr7 | 12860909 | 12866724 |
| chr7 | 18408502 | 18427426 |
| chr7 | 20241238 | 20250281 |
| chr7 | 46908925 | 46934128 |
| chr7 | 46969048 | 46983789 |
| chr7 | 47036365 | 47040369 |
| chr7 | 77281466 | 77284522 |
| chr7 | 77309525 | 77329065 |
| chr7 | 84022383 | 84061448 |
| chr7 | 84291523 | 84303271 |
| chr7 | 84477759 | 84492756 |
| chr7 | 112420418 | 112423266 |
| chr7 | 130868043 | 130928130 |
| chr7 | 130994541 | 131040253 |
| chr7 | 144820960 | 144828171 |
| chr8 | 32226850 | 32229337 |
| chr8 | 74985025 | 75004480 |
| chr8 | 77565552 | 77567476 |
| chr8 | 127870507 | 127890386 |
| chr8 | 128536531 | 128556866 |
| chr9 | 1707230 | 1708658 |
| chr9 | 3698128 | 3733512 |
| chr9 | 3906728 | 3951622 |

| | | |
|---|---|---|
| chr9 | 26243607 | 26262665 |
| chr9 | 73109166 | 73183814 |
| chr9 | 82488678 | 82489765 |
| chr9 | 82923315 | 82924813 |
| chrX | 45747751 | 45756995 |
